# The NK receptor NKp46 recognizes ecto-calreticulin on ER-stressed cells

**DOI:** 10.1101/2021.10.31.466654

**Authors:** Sumit Sen Santara, Ângela Crespo, Dian-Jang Lee, Jun Jacob Hu, Ying Zhang, Sourav Chowdhury, Karla F. Meza-Sosa, Marjorie Rowe, Arthur McClelland, Hao Wu, Caroline Junqueira, Judy Lieberman

**Affiliations:** Program in Cellular and Molecular Medicine, Boston Children’s Hospital, Boston MA USA; Department of Pediatrics, Harvard Medical School, Boston MA USA; Department of Biological Chemistry and Molecular Pharmacology, Harvard Medical School, Boston MA USA; Department of Chemistry and Chemical Biology, Harvard University, Cambridge MA, USA; Laboratorio de Neuroimmunobiología, Departamento de Medicina Molecular y Bioprocesos, Instituto de Biotecnología, Universidad Nacional Autónoma de México, Cuernavaca, Mexico; Center for Nanoscale Systems, Faculty of Arts and Sciences, Harvard University, Cambridge MA, USA; Instituto René Rachou, Fundação Oswaldo Cruz, Belo Horizonte, Brazil

## Abstract

Natural killer cells (NK) are a first line of immune defense to eliminate infected, transformed and stressed cells by releasing cytotoxic granules^1^. NK activation is controlled by the balance of signals transmitted by activating and inhibitory receptors but activating receptor engagement is required to trigger cytotoxicity. The activating receptor NKp46, encoded by the *NCR1* gene, is expressed by virtually all NK cells and is the most evolutionarily ancient NK receptor. NKp46 plays a major role in NK recognition of cancer cells, since NKp46 blocking antibodies potently inhibit NK killing of many cancer targets^2,3^. Although a few viral, fungal and soluble host ligands^4^ have been identified, the endogenous cell-surface ligand of this important activating NK receptor is unknown. Here we show that NKp46 recognizes and is activated by the P-domain of externalized calreticulin (ecto-CRT). CRT, normally localized to the ER, translocates to the cell surface during ER stress and is a hallmark of chemotherapy-treated dying cancer cells that induce an immune response (immunogenic cell death, ICD)^5^. NKp46 caps with ecto-CRT in NK immune synapses formed with ecto-CRT-bearing target cells. ER stress, induced by ZIKV infection, ICD-causing chemotherapy drugs and some senescence activators, externalizes CRT and triggers NKp46 signaling. NKp46-mediated killing is inhibited by *CRT* knockout or knockdown or anti-CRT antibodies and is enhanced by ectopic expression of GPI-anchored CRT. *NCR1/Ncr1*-deficient human and mouse NK are impaired in killing ZIKV-infected, ER-stressed, and senescent cells and cancer cells that endogenously or ectopically express ecto-CRT. Importantly, NKp46 recognition of ecto-CRT controls the growth of B16 melanoma and RAS-driven lung cancer in mouse models and enhances tumor-infiltrating NK degranulation and cytokine secretion. Thus, ecto-CRT is a danger-associated molecular pattern (DAMP) that is an endogenous NKp46 ligand that promotes innate immune elimination of ER-stressed cells.

---

NKp46 is constitutively expressed on virtually all NK and is also expressed on some innate lymphoid cells (ILC1/3)^6^. Mice deficient in *Ncr1*, the gene encoding NKp46, are impaired in tumor immune surveillance, have more severe influenza A, metapneumovirus, reovirus, *Candida glabrata*, pneumococcus and fusobacterium infections and more severe graft versus host disease^4,7,8^. The frequency of NKp46^+^ tumor-infiltrating lymphocytes (TIL) inversely correlates with metastatic disease of human gastrointestinal stromal tumors^9^. Despite the importance of NKp46 in NK immunity, the endogenous cell-surface ligand and activator of NKp46 is not known. The known ligands of NKp46/NCR1 are mostly pathogen products, such as viral hemagglutinins and fungal adhesins^10–12^ and soluble properdin, which doesn’t trigger classical NK activating receptor signaling^13^. Although NK recognize stressed and senescent cells, what types of stress are recognized and the stress and senescence ligands of NK activating receptors displayed on NK target cells are largely unknown^14^. The exceptions are the MIC/ULBP family ligands of NKG2D^15^ induced by some infections and genotoxic stress and soluble PDGF-DD for NKp44^16^.

CRT is a multifunctional, ER resident, Ca^++^-binding protein that acts as a sink for Ca^++^ to regulate cell signaling and as an unfolded protein chaperone^17^. ER stress and some forms of cell death cause CRT to traffic to the cell surface by an unclear mechanism to become ecto-CRT^5,18^. Ecto-CRT correlates with immunogenic cell death (ICD), namely cell death that elicits protective immunity^5^. Anti-tumor protection by ecto-CRT in ICD has been attributed to its function as an “eat-me” signal that triggers phagocytosis of ER-stressed and dying cells. Here we show that NKp46 recognizes stressed cells by binding to ecto-CRT, which is externalized in response to ER stress, but is constitutively expressed in some cancer cells and is induced by cytostatic drug-induced senescence, some viral infections, and treatment with some chemotherapeutic drugs.

## NK recognize and kill ZIKV-infected and ER-stressed cells

Human healthy donor peripheral blood NK degranulated and killed a ZIKV-infected, but not uninfected, human choriocarcinoma cell line, JEG-3 (**Fig. 1a,b**). Killing was mediated by cytotoxic granules, since it was inhibited by Ca^++^ chelation, which blocks granule exocytosis and perforin (PFN) activity but not death receptor-mediated killing (**data not shown**). ZIKV replicates in the ER and is known to induce ER stress^19^, which we verified by qRT-PCR assessment of *BIP*, *CHOP* and spliced *XBP1* mRNA (**Fig. 1c**). ZIKV induction of these mRNAs was comparable to that induced by the ER stressor tunicamycin (Tu) and was inhibited by the ER-stress inhibitor salubrinal, which inhibits dephosphorylation of the eukaryotic translation initiation factor 2α (eIF2α)^20^. NK do not degranulate or kill CMV^21^ or HSV-2 infected JEG-3. ER stress is not activated by these human herpesviruses^22^, which we confirmed in JEG-3 (**Fig. 1c**). Based on these data, we hypothesized that NK killing of ZIKV-infected cells might be mediated by recognizing some feature of ER stress. To test this hypothesis, we compared peripheral blood NK killing of ZIKV-infected or Tu-treated JEG-3 (**Fig. 1d**). Both ER stress inducers activated NK killing, which was blocked to baseline levels by salubrinal. NK killing of ZIKV-infected JEG-3 was significantly inhibited to different extents by ER stress inhibitors - most strongly by the S1P inhibitor PF-429242, which inhibits the ATF6 pathway, and weakly by the PERK inhibitor GSK2606414 but not significantly by the IRE1α inhibitor 4μ8C or by TUDCA (tauroursodeoxycholic acid), a bile acid that acts as an ER chaperone (**Extended Data Fig. 1a**). Thus, human NK are triggered to kill by recognizing some feature of ER stress.

**Figure 1.**
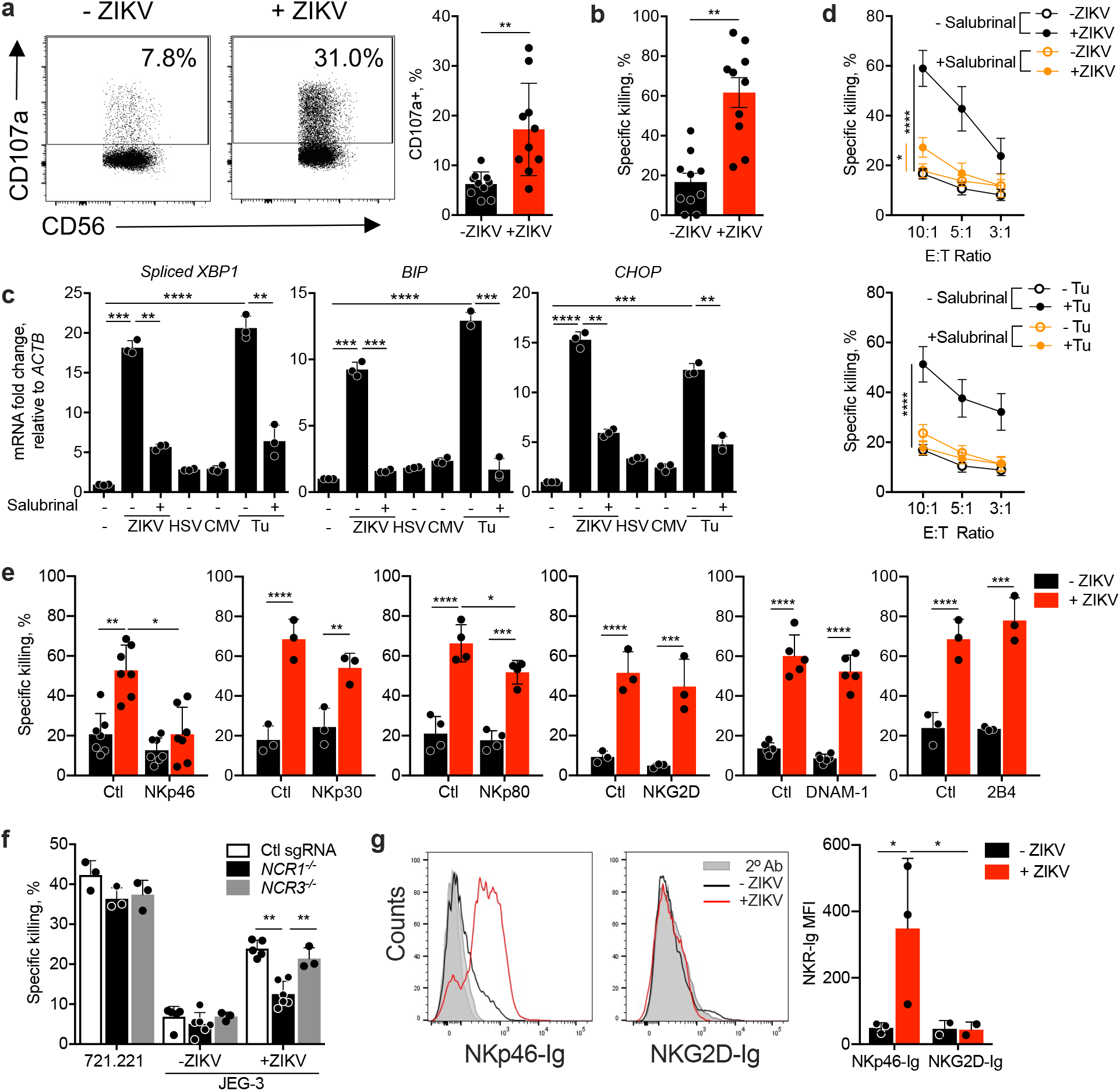
NK recognize ZIKV-infected and ER-stressed target cells. **a,** Representative flow cytometry plots (left) and percentage of degranulating NK, isolated from the blood of healthy donors (n=10, right), measured by surface CD107a, in response to uninfected and ZIKV-infected JEG-3 (8 h co-culture, E:T ratio 1:3). **b,** NK specific killing of uninfected and ZIKV-infected JEG-3. **c,** ER stress, assessed by *XBP1* splicing (left) and increases in *BIP* (middle) and *CHOP* (right) mRNA, in JEG-3 that were uninfected or infected with ZIKV, HSV-2 or HCMV for 1-2 d or treated with tunicamycin (Tu) for 1 d. Indicated samples were pretreated with the ER stress inhibitor salubrinal (n=6). mRNA levels assayed by qRT-PCR, were normalized to *ACTB*. **d,** Effect of salubrinal pretreatment of target cells on NK killing of ZIKV-infected (top) or Tu-treated (bottom) JEG-3 (n=6). **e,** Effect of NK receptor blocking antibodies on NK killing of uninfected or ZIKV-infected JEG3 (n=3-7). **f,** Specific killing of the classical NK target 722.221 or of uninfected or ZIKV-infected JEG-3 by the human NK cell line YT knocked out for *NCR1* or *NCR3* or treated with control sgRNAs (n=3-6). **g,** Representative flow cytometry histogram (left) and mean fluorescence intensity (MFI) of NK receptor (NKR)-Ig fusion protein (NKp46-Ig and NKG2D-Ig) binding to uninfected or ZIKV-infected JEG-3 (right) (n=3). Specific killing in (b,d-f) assessed by 8h ^51^Cr release assay using an E:T ratio of 10:1 unless otherwise indicated. Graphs show mean ± SEM of at least 3 independent experiments or technical replicates. Statistics were performed using two-tailed non-parametric unpaired *t*-test (a,b), one-way ANOVA (c,e-g), or area under the curve, followed by one-way ANOVA (d). *P*: *<0.05, **<0.01, ***<0.001, ****<0.0001.

## NKp46 binds to ZIKV-infected cells and mediates NK killing

To identify the activating receptor responsible for NK killing of ZIKV-infected JEG-3, purified healthy donor peripheral blood NK killing was assessed in the presence or absence of blocking antibodies to widely expressed NK activating receptors (**Fig. 1e**). Blocking NKp46 strongly and significantly inhibited NK killing, suggesting that NK use NKp46 to recognize ZIKV-infected cells. Blocking NKp80 had a weak, but significant effect on killing of ZIKV-infected cells but blocking antibodies to other NK activating receptors (NKp30, NKG2D, DNAM-1, 2B4) had no significant effect. We therefore focused on NKp46. To confirm a role for NKp46 in killing ZIKV-infected cells, CRISPR/Cas9 was used to knockout *NCR1*, the gene encoding NKp46, or *NCR3*, the gene encoding NKp30, in the human NK cell line YT (**Extended Data Fig. 1b,c**; subsequent experiments were performed with at least two different KO clones, but results for only one clone are shown). Although knockout of *NCR1* or *NCR3* in YT did not significantly affect killing of the classical NK target cell 721.221, *NCR1^−/−^* YT killing of ZIKV-infected JEG-3 was reduced ~2-fold compared to killing of YT transfected with control sgRNAs. Killing by *NCR3^−/−^* YT was not significantly changed (**Fig. 1f**). Next, to verify a role of NKp46 in recognizing ZIKV-infected JEG-3, uninfected or ZIKV-infected JEG-3 binding to the extracellular domains of NKp46 or NKG2D fused to the Fc region of human IgG1 (NKp46-Ig, NKG2D-Ig) was assessed by flow cytometry (**Fig. 1g**). (We verified that NKp46-Ig did not aggregate by size exclusion chromatography, which can give spurious results (**Extended Data Fig. 1d**).) NKp46-Ig, but not NKG2D-Ig, selectively bound to ZIKV-infected JEG-3. Thus, NKp46 recognizes ZIKV-infected cells and activates NK killing.

## NKp46 binds to ecto-calreticulin

To identify putative NKp46 ligands, NKp46-Ig and NKG2D-Ig were used to pull down cross-linked potential ligands on the ZIKV-infected JEG-3 cell membrane (**Fig. 2a**, **Extended Data Fig. 1a**). A single dominant high molecular weight NKp46-Ig band in ZIKV-infected cells was identified by silver staining, but no extra band was detected in the NKG2D-Ig pulldown. Proteins in the NKp46-Ig pulled down band were identified by mass spectrometry as the ER chaperones, calreticulin (CRT) and protein disulfide isomerases (PDI, PDIA1 and PDIA3). (Two lower molecular weight bands in the NKp46-Ig pulldown did not contain cross-linked fusion protein but contained fragments of protein A and BSA, which were judged contaminants.) The CRT and PDI ER chaperones selectively translocate and associate on the cell surface during ER stress through a poorly understood mechanism, while other ER chaperones, such as calnexin and Hsp70, are not externalized^18,23,24^. Cell surface staining of CRT and PDI increased with ZIKV infection or Tu, but Hsp70 staining did not change (**Fig. 2b**). To test whether NKp46 senses ecto-CRT and/or PDI, we took advantage of the fact that untreated JEG-3 express little ecto-CRT, but the ICD-causing chemotherapy drug oxaliplatin (OP) induces ER stress and CRT externalization, which was verified (**Fig. 2c**). NKp46-Ig pulled down CRT from OP-treated, but not untreated, JEG-3, but NKp44-Ig or NKp30-Ig did not pulldown CRT from either (**Fig. 2d**). As a specificity control, HLA-G, expressed on the surface of JEG-3, was not pulled down by either fusion protein. Thus, NKp46 specifically binds to ecto-CRT.

**Figure 2.**
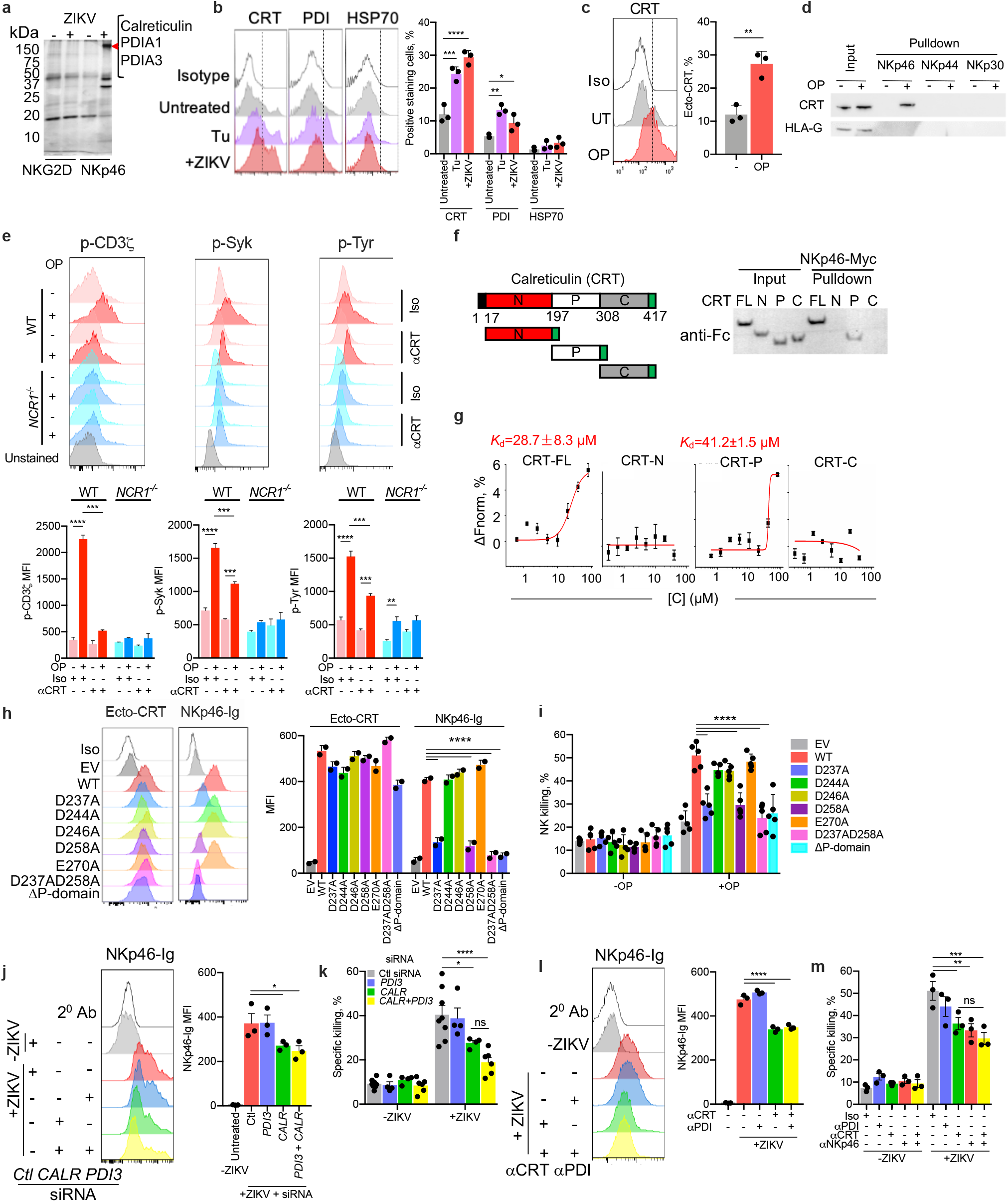
NKp46 recognizes the P domain of ecto-calreticulin. **a,** Immunoblot of proteins pulled down and chemically crosslinked with NKG2D-Ig and NKp46-Ig. The high molecular weight band in ZIKV-infected cells (red arrow) was analyzed by mass spectrometry for candidate NKp46 ligands. The proteins with the highest peptide coverage are indicated at right. Mass spectrometry of the NKp46-Ig low molecular weight bands specific for ZIKV-infected cells did not contain NKp46-fusion protein peptides but contained peptide sequences of fragments of contaminating BSA and protein A. **b,** Representative flow cytometry histograms of surface CRT, PDI and HSP70 on uninfected, tunicamycin-treated (Tu) or ZIKV-infected JEG-3 (left) and fluorescence intensity of 3 independent samples (MFI, right). **c,** Flow cytometry histograms of surface CRT expression on JEG-3 that were untreated or treated with oxaliplatin (OP) (left) and MFI of 3 samples (right). **d,** CRT immunoblot of proteins pulled down with NKp46-Ig, NKp44-Ig and NKp30-Ig from the membrane protein-enriched faction of JEG-3 treated or not with OP. Immunoblot was probed for HLA-G as control. **e,** Phospho-flow cytometry of YT (WT or knocked out using CRISPR/Cas9 for *NCR1*) incubated for 15 min with OP-treated or untreated JEG-3 in the presence or absence of anti-CRT or control antibody and co-stained for CD56 and p-CD3ζ (left), p-Syk (middle) and p-Tyr (right). YT was preincubated with anti-NKG2D to reduce background signaling from this NK receptor. Top shows representative histograms and bottom shows fluorescence intensity of 2 independent samples and statistical comparison of signaling in samples that used WT or *NCR1^−/−^* YT. **f,** (top) Schematic of CRT domains - signal peptide (black); N-domain (red); P-domain (white); C-domain (grey). Full-length (FL) or individual CRT domains were fused with a C-terminal Fc tag (green). (bottom) CRT-Fc fusion proteins were incubated with Myc-tagged NKp46 and pulled down with anti-Myc beads. Samples were probed with anti-Fc. **g,** Binding of Alexa647-labeled NKp46-Myc to Fc-tagged FL and N, P, or C domains of CRT was analyzed by microscale thermophoresis (dissociation constants of FL and P domain are shown). **h,** Representative flow cytometry histograms of CRT surface expression and NKp46 binding (left) and MFI (right) of 2 individual samples of *CALR* knockout HEK293T cells transfected with empty vector (EV) or WT or mutated CRT and treated with OP. **i,** Killing by human primary blood NK of 293T cells transfected as in (**h**) with empty vector (EV) or WT or mutated CRT and treated with OP (4 h ^51^Cr release, E:T ratio 10:1, n=5). **j,** JEG-3, knocked down for *CALR* and/or *PDI3 or* with nontargeting (Ctl) siRNAs were infected or not with ZIKV and analyzed for NKp46-Ig binding by flow cytometry (left); NKp46-Ig MFI of untreated or knocked down cells in 3 samples (right). **k,** Healthy donor NK killing by ^51^Cr release assay (4 h, E:T ratio 10:1, n=6) against knocked down JEG-3. **l,** Effect of anti-CRT and/or anti-PDI on NKp46-Ig binding to JEG-3 that were infected or not with ZIKV. Shown are representative flow histograms (left) and MFI of 3 samples (right). **m,** Effect of anti-CRT, anti-PDI and anti-NKp46 on healthy donor NK killing of uninfected or ZIKV-infected JEG-3 (4 h, E:T ratio 10:1, n=3). Data are representative of three independent experiments. Statistics were performed using one-way ANOVA (b,h-m), parametric unpaired *t*-test (c) and two-way ANOVA (e). *P*: *<0.05, **<0.01, ***<0.001, ****<0.0001.

## NKp46 binding to ecto-CRT activates NK signaling

To determine whether NKp46 binding to ecto-CRT leads to NK activation, phosphorylation of the CD3ζ NKp46 adaptor, downstream Syk protein tyrosine kinase and overall p-Tyr was measured after WT and *NCR1^−/−^* YT were incubated with with untreated or OP-treated JEG-3 in the presence of blocking antibody to NKG2D to inhibit triggering of this NKR (**Fig. 2e**). In WT YT exposed to OP-treated JEG-3, phosphorylation of CD3ζ, Syk or Tyr were all increased and phosphorylation was inhibited by anti-CRT. Moreover, OP treatment of JEG-3 did not increase CD3ζ or Syk phosphorylation of *NCR1^−/−^* YT when NKG2D signaling was blocked. Thus, NKp46 sensing of ecto-CRT activates NK signaling.

## NKp46 binds to the P domain of ecto-calreticulin

CRT has 3 domains – an N terminal globular domain with chaperone function, a proline-rich P domain that has lectin properties and is responsible for most protein interactions, and a C terminal acidic domain that binds Ca^++^ and has an ER retention signal^25^ (**Fig. 2f**). To determine which domain is recognized by NKp46, full length and individual domains of CRT fused at the C-terminus to the Fc domain of human Ig were incubated with Myc-tagged NKp46 (**Fig. 2f**). When NKp46 was pulled down from the incubated proteins and immunoblotted with anti-Fc, both full-length (FL) and to a somewhat lesser extent, the CRT P domain, but not the N or C domains, were detected. Microscale thermophoresis (MST) was used to quantify the strength of the NKp46-CRT interaction, (**Fig. 2g**). Consistent with the pulldown, the Myc-tagged extracellular domain of NKp46 bound to Fc-tagged full length and P domain CRT, but not N or C CRT, with respective K_d_s of 28.7 ± 8.3 μM and 41.2 ± 1.46 μM. Thus, NKp46 binds to the P domain of exposed CRT. The K_d_ measured by MST for binding to His-tagged CRT was 12.3 ± 2.4 μM for NKp46-Ig, ~5-50-fold weaker than for other reported ligands of NKp46 (hemagluttinin (HA), CD4, Siglec6 and Siglec8) (**Extended Data Fig. 2b,c**). Although P domain binding to some ligands, such as influenza HA, depends on NKp46 sialylation, neuraminidase treatment of NKp46 did not significantly affect CRT binding. Raman spectroscopy confirmed NKp46 binding to CRT (**Extended Data Fig. 2d,e**). A unique concentration-dependent peak at 1658 cm^−1^ in the Raman spectrum of the amide I region was detected in the CRT and NKp46 mixture that was absent in the spectrum of either protein on its own. Hyperbolic fitting of the peak area vs CRT concentration provided an estimate of 9.95 ± 5 μM for the K_d_ of the interaction, similar to the K_d_s obtained by MST. The measured NKp46-CRT K_d_ values are on the high end of the range of ligand binding constants of other NK receptors, but within the range of T cell receptor K_d_s^26–28^.

The ectodomain of NKp46 consists of 2 immunoglobulin domains connected by an interdomain hinge region (**Extended Data Fig. 3a**)^29^. In homologous receptors, ligand binding is mediated by charge interactions in the hinge region or by the exposed end of the first immunoglobulin domain. The P domains of CRT and other structurally related lectin chaperones bind to a variety of proteins, including ER folding factors, by interactions of the negatively charged tip of a hairpin formed by proline-rich modules (**Extended Data Fig. 3b**). Residues M257 and D258 at the tip of the P domain within a conserved sequence that is rich in prolines and charged residues play an important role in P domain interactions^25^. We used CLUSPro and the PIPER algorithm to model the interaction of CRT binding onto the human NKp46 ectodomain^30^ (**Extended Data Fig. 3c-e**). Docking suggested that the CRT P domain inserts into a channel in the hinge region and is grasped between the 2 Ig-like domains of NKp46. Conserved R160 and K170 on NKp46, located at opposite extremes of the cleft, form the P-domain binding groove. In the modeled docked complex, positively charged residues on NKp46 (R160, K170, R166, and H197) form salt bridges with D258, S231, D237 and E270, respectively, of the P-domain (**Extended Data Fig. 3d,e**). The predicted binding energy (ΔG, −10kcal mol-1 at 298° K) and dissociation constant (K_d_=19.9±3.4 μM), calculated from an ensemble of bound/docked conformations, suggests a potentially stable docked complex and also is consistent with the MST and Raman binding constants (**Fig. 2g**, **Extended Data Fig. 2b-e**).

## Knocking down *CALR* and anti-CRT inhibit NKp46-Ig binding and NK killing

To confirm CRT as an NKp46 ligand and identify the amino acid residues important for binding, we used CRISPR/Cas9 to knock out *CALR*, the gene encoding CRT, in HEK293T and rescued CRT expression with WT *CALR* or *CALR* encoding for Ala mutations of conserved acidic residues in the tip of the P domain (D237, D244, D246, D258, and E270) or deleted of the P-domain (ΔP) **(Extended Data Fig. 4a)**. After OP treatment, ecto-CRT was not detected in *CALR^−/−^* HEK293T cells but was detected at comparable levels for all the rescued constructs (**Fig. 2h**). NKp46-Ig did not bind above background to OP-treated *CALR^−/−^* HEK293T, but NKp46-Ig binding was rescued by WT, D244A, D246A, or E279A *CALR*. Binding was not rescued by ΔP, D237A or D258 or the double D237AD258A mutant *CALR*, confirming the identification of the P-domain as the binding site and our modeling of the interaction between the tip of the P-domain and the NKp46 hinge domain. NK did not kill OP-treated *CALR^−/−^* HEK293T but did kill all the knockout cells rescued with mutations that maintained NKp46-Ig binding, but none of the cells rescued with binding-deficient CRT mutants (**Fig. 2i**). Thus, D237 and D258 in the P-domain of CRT mediate binding to NKp46 and NK killing of target cells expressing ecto-CRT.

To further confirm ecto-CRT as an NKp46 ligand, JEG-3 cells were knocked down for *CALR* and/or *PDIA3* and NKp46-Ig binding and NK killing of knocked down ZIKV-infected JEG-3 was compared to infected JEG-3 treated with control nontargeting siRNA (**Fig. 2j,k**, **Extended Data Fig. 4b-d**). *CALR* and *PDIA3* knockdown did not change surface MHC-I expression on uninfected JEG-3 cells, but both significantly reduced surface CRT on ZIKV-infected JEG-3 cells. The effect of *PDIA3* knockdown on ecto-CRT was modest (~20%) compared to *CALR* knockdown (~80%) and was not unexpected as PDI is known to facilitate the exposure of CRT after ER stress^25^. Knockdown of *CALR*, but not *PDIA3*, significantly reduced NKp46-Ig binding to ZIKV-infected cells and NK killing of ZIKV-infected JEG-3 (**Fig. 2j,k**). Knocking down both *CALR* and *PDIA3* was not significantly different from knocking down just *CALR*.

The importance of ecto-CRT in binding NKp46 and triggering killing was also evaluated by comparing NKp46-Ig binding and peripheral blood NK killing of ZIKV-infected JEG-3 in the presence of antibodies to CRT, NKp46, PDI or an isotype control antibody (**Fig. 2l,m**). Like the knockdown, anti-CRT, but not anti-PDI, significantly reduced both NKp46-Ig binding and NK killing of ZIKV-infected JEG-3. Anti-PDI did not inhibit NK killing of ZIKV-infected JEG-3 but anti-CRT and anti-NKp46 both similarly inhibited killing. Combining anti-CRT and anti-NKp46 did not significantly increase the inhibition of either antibody on its own, consistent with our hypothesis that ecto-CRT and NKp46 operate in the same pathway by binding to each other. Thus, NKp46 binds to ecto-CRT to trigger killing of target cells with exposed CRT.

## NCR1 recognizes ecto-CRT on mouse tumor lines treated with an ICD-inducing drug

NCR1/NKp46 is the only NCR expressed in most mouse strains. To determine whether mouse NCR1 also recognizes ER-stressed cells by binding to exposed CRT, B16-F10 melanoma (hereafter referred to as B16) that were untreated or treated with cisplatin (CP), which is not an ICD drug, or OP, which is an ICD drug,^31^ were analyzed for ecto-CRT by flow cytometry. Both untreated and CP-treated B16 showed minimal ecto-CRT above background, while ecto-CRT was strongly induced by noncytotoxic concentrations of OP (**Fig. 3a**). To determine whether NK killing of OP-treated B16 depends on NCR1 recognition of ecto-CRT, mouse splenic NK killing of untreated, OP or CP-treated B16 was assessed in the presence or absence of blocking antibody to CRT or NCR1 (**Fig. 3b**). NK killing of OP and CP-treated B16 was enhanced about 2-fold compared to untreated cells. Both these drugs upregulate NKG2D stress ligands^32,33^. The small amount of baseline killing of untreated B16 was significantly and similarly inhibited by antibodies to CRT or NCR1. Blocking CRT or NCR1 strongly inhibited killing of OP-treated B16 to a similar extent but had no significant effect on killing of CP-treated B16, which did not expose ecto-CRT. Consistently, low level killing of untreated B16 by *Ncr1^−/−^* splenic NK, but not by *Klrk1^−/−^* NK, was significantly reduced. More importantly, killing of OP-treated B16 by splenic NK from *Ncr1^−/−^* mice was strongly reduced compared to killing by wild-type (WT) splenic NK, but was not significantly impaired in killing CP-treated B16 (**Fig. 3c**). In contrast, splenic NK isolated from *Klrk1^−/−^* mice, deficient in NKG2D, significantly inhibited CP-treated B16 killing, but had no significant effect on killing OP-treated cells. These results taken together indicate that NCR1 recognition of ecto-CRT contributed to killing of OP-treated, but not CP-treated, B16 (**Fig. 3c**).

**Figure 3.**
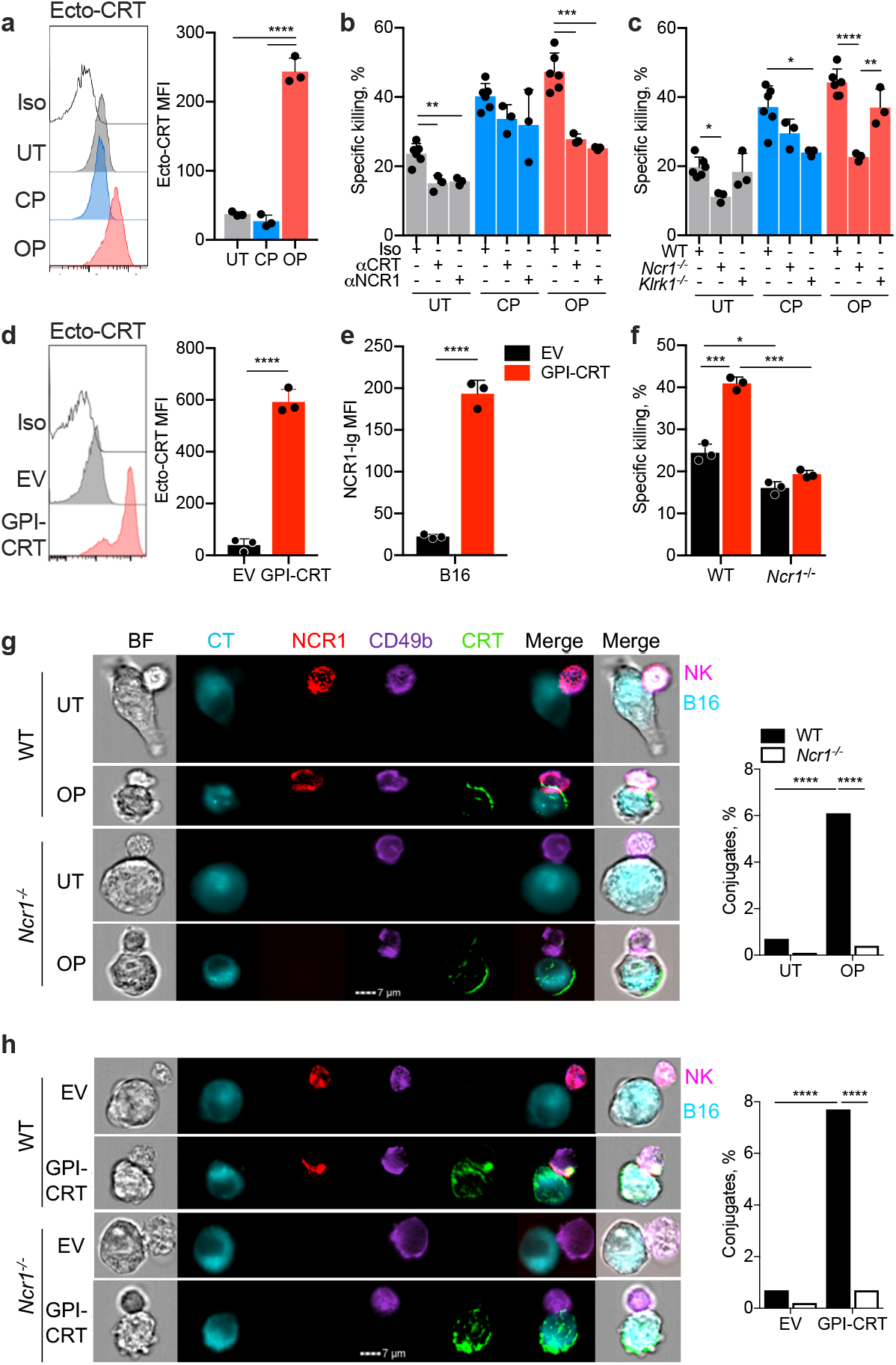
NKp46 recognition of ecto-CRT enhances immune synapse formation and NK killing of tumor cell lines. **a,** Representative flow cytometry histograms of CRT surface expression (left) and mean±SEM ecto-CRT mean fluorescence intensity (MFI, right) of untreated (UT), oxaliplatin (OP) or cisplatin (CP) treated B16. Iso, staining with isotype control Ab (n=3). **b,** Effect of anti-CRT on mouse splenic NK killing of UT or OP- or CP-treated B16 in the presence of isotype control antibody, anti-CRT or anti-NCR1 (n=3 or 5). **c,** Killing by splenic NK from WT, *Ncr1^−/−^* or *Klrk1^−/−^* mice of B16 that were UT or treated with CP or OP (n=3 or 5). **d,** Representative flow cytometry histograms of CRT surface expression (left) and ecto-CRT MFI of 3 samples (right) of B16 stably transfected with EV or to express GPI-linked CRT. **e,** Effect of ectopic ecto-CRT on NCR1-Ig binding to B16. **f,** WT or *Ncr1^−/−^* splenic NK cell killing of EV or GPI-CRT overexpressing B16 (n=3). Killing in (b,c,f) assessed by 4 h ^51^Cr release, E:T ratio 10:1. **g,h,** Imaging flow cytometry analysis of conjugates of splenic NK isolated from WT or *Ncr1^−/−^* mice, as indicated, with B16 that were untreated or treated with OP **(g)** or transfected with empty vector (EV) or GPI-CRT plasmid **(h).** Shown are representative images of conjugates (left) and the proportion of cells forming NK:target cell conjugates (right). Cells were stained with CellTracker Red CMTPX (blue), anti-NCR1 (red), anti-CD49b (violet) and anti-CRT (green). Statistics in (g,h) were performed using a χ^2^ test. Data shown are representative of ~1000 individual images. Shown images are representative of at least 3 independent experiments. Statistics by unpaired one-way ANOVA (a-c), parametric unpaired *t*-test (d,e), two-way ANOVA (f) and *Chi*-square test (g,h). *P*: *<0.05; **<0.01; ***<0.001; ****<0.0001.

## Engineered ecto-CRT over-expression enhances NKp46 activation and NK killing

To confirm that ecto-CRT triggers NKp46, B16 were transfected with an expression plasmid encoding CRT with a C-terminal glycophosphatidylinositol (GPI) anchor to generate a stable B16-GPI-CRT line^34^. Cell surface CRT expression on engineered B16-GPI-CRT increased dramatically compared to cells transfected with the empty vector (EV), but was only about 2-fold higher than OP-induced endogenous levels (**Fig. 3d**). Even though CRT is a member of a multi-protein peptide-loading complex, which is involved in MHC class I folding and peptide loading for antigen presentation, ectopic ecto-CRT had no effect on class I MHC cell surface expression, as assayed by flow cytometry staining of H-2K^b^ (**Extended Data Fig. 5a**). Ectopic ecto-CRT also had no effect on in vitro proliferation of either cell line and slightly, but significantly, increased colony formation (**Extended Data Fig. 5b,c)**. Compared with WT or EV B16, B16-GPI-CRT showed significantly increased cell migration and invasion in Transwell assays (**Extended Data Fig. 5d,e**). These in vitro experiments suggest that ecto-CRT expression might increase intrinsic B16 malignant properties.

Enhanced expression of ecto-CRT significantly increased binding of NCR1-Ig to engineered B16 (**Fig. 3e**). Consistently, WT splenic NK killing of the engineered cell line expressing ectopic ecto-CRT was increased compared to EV-transfected cells but there was no increased killing by splenic NK from *Ncr1^−/−^* mice (**Fig. 3f**), indicating that ecto-CRT recognition and NK activation in mice was mediated solely by NCR1.

The P domain of CRT is highly conserved in mouse and humans (97.3% identity with perfect conservation of the region modeled to bind to NKp46; **Extended Data Fig. 3f**). To confirm CRT as an NCR1 ligand and test whether the CRT P domain interacts with NCR1 as to human NKp46, WT and mutant (D237A, D244A, D258A, ΔP) GPI-anchored mouse CRT was expressed in B16 and cells were assayed for ecto-CRT, NCR1-Ig binding and susceptibility to splenic NK killing (**Extended Data Fig. 3g,h**). Although all the mutants expressed CRT similarly, NCR1-Fc binding increased only to cells expressing WT or D244A CRT, but not to cells expressing CRT containing the critical mutations that abrogated binding to human NKp46. Consistently, NK killing increased above background only for targets expressing WT or D244A GPI-CRT. Thus, in mice, as in humans, conserved Asp residues in the CRT P domain control NCR1 binding and activation.

## NKp46 and calreticulin co-localize at the immune synapse

When NK activating receptors recognize a target cell, the NK and its target cell form an immunological synapse (IS) where the activating NK receptors and their ligands cap^35^. To confirm that ecto-CRT is the ligand of NKp46, the number of splenic NK conjugates formed with B16 that were treated or not with OP or were stably transfected with GPI-CRT or EV plasmids was quantified by imaging flow cytometry (**Fig. 3g,h**). The number of NK-B16 conjugates increased about 10-fold after OP treatment or GPI-CRT expression. Moreover, images of the conjugates formed with these ecto-CRT^+^ B16 showed capping of NKp46 and the integrin CD49b on the NK with CRT on the target cell, whereas no such capping was visualized in the few conjugates formed with B16 that do not display ecto-CRT. When *Ncr1^−/−^* splenic NK were substituted for WT NK, the number of conjugates formed with OP-treated B16 or B16-GPI-CRT dropped to baseline levels and no capping was observed. These data further confirm that NKp46 recognizes ecto-CRT.

## NCR1 recognition of ecto-calreticulin inhibits B16 tumor growth in mice

To examine the role of NCR1 recognition of ecto-CRT in anti-tumor immunity, WT and *Ncr1^−/−^* mice were inoculated sc with B16-EV or B16-GPI-CRT (**Fig. 4a-c, Extended Data Fig. 6**). Although B16-EV tumors grew similarly in WT and *Ncr1^−/−^* mice, B16-GPI-CRT growth was suppressed in WT mice, but not in *Ncr1^−/−^* mice (**Fig. 4a**). In fact, B16-GPI-CRT tumors unexpectedly grew more rapidly than B16-EV tumors in *Ncr1^−/−^* mice, perhaps because ecto-CRT increased in vivo malignant properties as it did in vitro (**Extended Data Fig. 5c-e**). Thus, NCR1 suppressed the growth of ecto-CRT-expressing B16 in WT mice. To identify immune differences in tumor infiltrating cells that might account for the protection afforded by ecto-CRT expression, CD45^+^ cells were isolated from B16-EV or B16-GPI-CRT tumors 3 weeks after implantation in WT and *Ncr1^−/−^* mice. Although the numbers of tumor-infiltrating NK, CD8 T lymphocytes, dendritic cells (DC) or tumor-associated macrophages (TAM) did not significantly differ in the four groups of tumors (**Extended Data Fig. 6a**), more tumor-infiltrating NK in WT mice bearing B16-GPI-CRT tumors stained for the cytotoxic effector proteins granzyme B (GzmB) and PFN than in *Ncr1^−/−^* mice (**Fig. 4b**). Similarly, the tumor-infiltrating NK cytokine secretion and degranulation functions in B16-GPI-CRT tumor-bearing mice were significantly increased in WT compared to *Ncr1^−/−^* mice (**Fig. 4c**). In contrast, GzmB and PFN staining in CD8 T cells was low but significantly greater in *Ncr1^−/−^* mice than WT mice and IFNγ production by CD8 TIL was significantly reduced in tumors from *Ncr1^−/−^* mice. These differences may be secondary to the modulating effect of NK responses on antigen-specific CD8 T cell functions^36^. Ecto-CRT acts as an “eat-me” signal that stimulates phagocytosis by DC or TAM. Since the B16-EV and B16-GPI-CRT cells used in this manuscript are DsRed^+^, DsRed staining of DC and TAM could be used to measure in vivo phagocytosis. WT tumor-infiltrating DC and TAM were not significantly more DsRed^+^ than the same cells from *Ncr1^−/−^* mice, suggesting enhanced phagocytosis may not have been the major factor responsible for better tumor control (**Extended Data Fig. 6b**). Thus, ectopic ecto-CRT expression on B16 enhances tumor-infiltrating NK functions in WT compared to *Ncr1^−/−^* mice, confirming the in vivo importance of NCR1 recognition of ecto-CRT.

**Figure 4.**
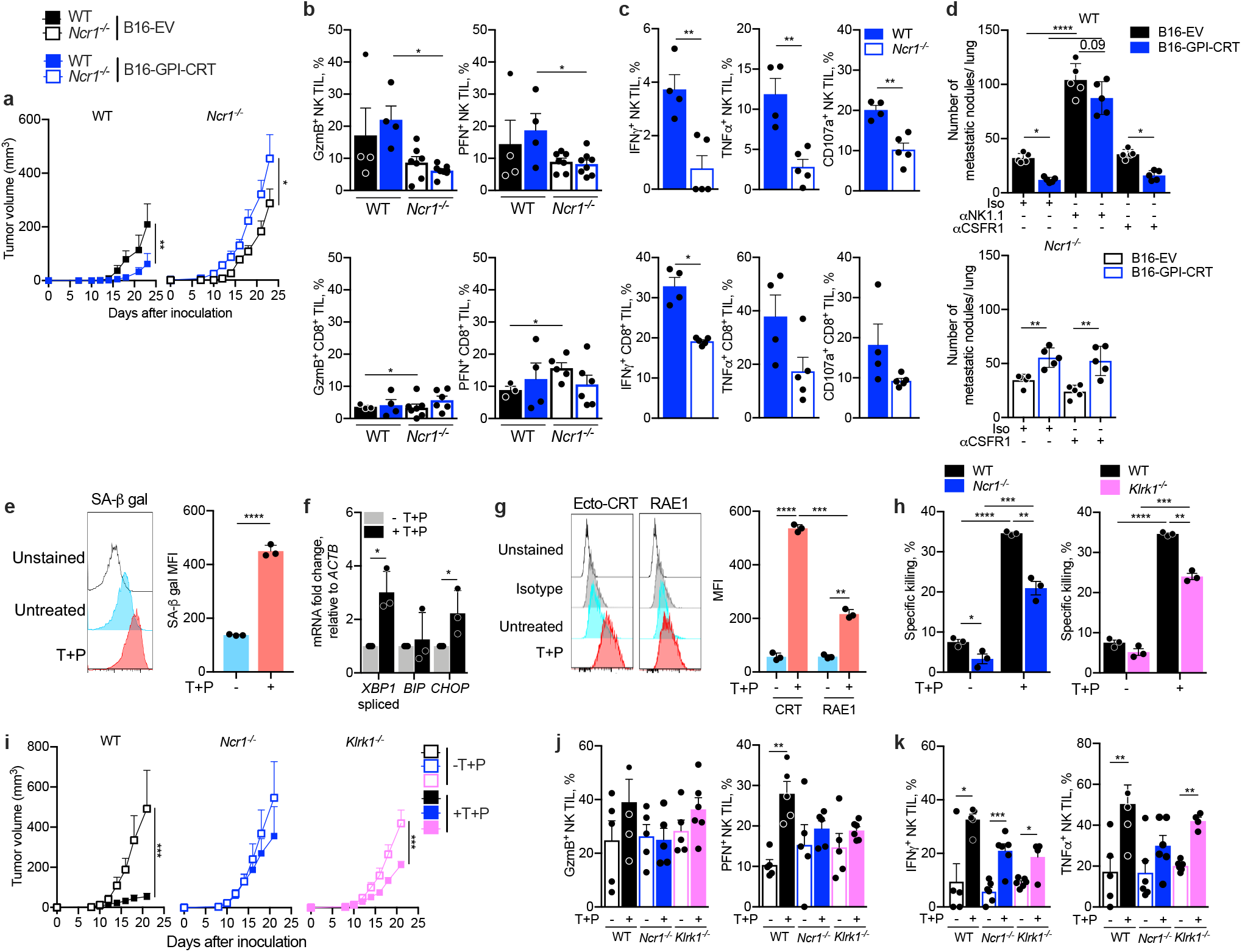
NKp46 recognition of ecto-calreticulin improves NK tumor control and antitumor immunity in mice and is activated during trametinib and palbociclib-induced senescence. **a-c,** Empty vector (EV) or GPI-CRT-overexpressing B16 were injected sc into WT or *Ncr1^−/−^* mice (n=7/group). Tumor growth (**a**) and functional markers (**b,c**) of NK (top) and CD8^+^ (bottom) TIL were assessed at time of sacrifice 24 days after tumor implantation (**b).** In **c**, PMA + ionomycin stimulated IFNγ (left) and TNFα (middle) production and degranulation, assessed by CD107a externalization (right) are shown. **d,** WT and *Ncr1^−/−^* mice were injected with 2 × 10^5^ empty vector (EV) or GPI-CRT expressing B16 via tail vein on day 0 into WT (top) or *Ncr1^−/−^* mice (bottom) (n=5). In indicated samples, NK cell depleting antibody (anti-NK1.1) was injected (i.p.) on days −1, 0 and once a week or anti-CSF1R was injected every other day to deplete macrophages. Tumor metastasis was assessed on day 20 by counting visible metastatic nodules on the lung surface. **d-f,** Trametinib and palbociclib (T+P) in vitro treatment of mouse KP lung cancer cells induces senescence and activates NK cell surface expression of NK activating ligands. Shown are (**e**) representative flow cytometry histograms of β-galactosidase activity (SA-β-gal, left) and MFI quantification of multiple samples (right), (**f**) ER stress, assessed by qRT-PCR of *Xbp1* splicing and *Bip* and *Chop* mRNA, and (**g**) CRT and RAE1 expression in mouse KP that were untreated or treated with T+P. (n=3) **h,** In vitro killing by splenic NK from WT, *Ncr1^−/−^* or *Klrk1^−/−^* mice of KP that were untreated or treated with T+P (4 h ^51^Cr release, E:T ratio 20:1). **i,j,** KP cells were injected sc into WT, *Ncr1^−/−^* or *Klrk1^−/−^* mice (n=5-7/group) and animals were treated with T+P by oral gavage 13-18 d after tumor implantation. Tumor growth (**i**) and NK TIL were assessed at time of sacrifice for cytotoxic granule proteins (**j**) and cytokine production (**k**) were assessed. Data in bar graphs show mean ± SEM of at least 3 independent experiments. Statistics calculated by area under the curve, followed by two-tailed nonparametric unpaired *t*-test (a,i), one-way ANOVA (b,d,g,h), two-tailed nonparametric unpaired *t*-test (c,e,j,k). *P*: *<0.05, **<0.01, ***<0.001, ****<0.0001

To determine whether NCR1-mediated protection involves CD8 T cells, WT and *Ncr1^−/−^* mice were depleted of CD8 T cells or treated with isotype control antibody before implanting B16-EV or B16-GPI-CRT tumors (**Extended Data Fig. 6c,d**). In WT mice ecto-CRT-expressing tumors grew significantly more slowly than EV B16 even when CD8 T cells were depleted (**Extended Data Fig. 6e**). In antibody-treated *Ncr1^−/−^* mice, B16-GPI-CRT tumors still grew more than B16-EV tumors but the difference was no longer significant in CD8-depleted mice. This depletion experiment suggests that protection afforded by NCR1 recognition of ecto-CRT does not depend on CD8 T cells, which may not be surprising since CD8 T cells do not express NCR1.

## Ecto-CRT expression on B16 reduces metastases and is NCR1 dependent

To investigate whether NCR1 recognition of ecto-CRT has a role in controlling metastasis, B16-EV and B16-GPI-CRT were injected into the tail vein of WT or *Ncr1^−/−^* mice and metastatic lung nodules were counted 3-4 weeks later (**Fig. 4c**). To assess the role of NK vs CRT-mediated phagocytosis in CRT-dependent protection, these mice were also depleted of NK or macrophages by injection of anti-NK1.1 or anti-CSFR1, respectively, or were injected with an isotype control antibody (**Extended Data Fig. 6c**). In undepleted or macrophage-depleted WT mice, the number of B16-GPI-CRT metastatic nodules was significantly lower than B16-EV nodules, but the number of nodules increased greatly (~2.5-fold) and GPI-CRT expression no longer significantly conferred protection in NK-depleted mice, indicating that NK were responsible for suppressing metastasis of tumors with exposed CRT (**Fig. 4c**, **top**). Depletion of macrophages did not significantly change the number of lung metastases for either tumor. Thus, ecto-CRT protection from metastases of B16 tumors is mediated by NK NCR1, with no strong role of macrophages. By contrast, ectopic expression of ecto-CRT in B16 tumors increased the number of lung metastases in *Ncr1^−/−^* mice, as it did for primary B16 tumors, and the number of metastases did not change with macrophage depletion (**Fig. 4c**, **bottom**). These results in both primary and metastatic models suggest that macrophages do not play a major role in enhanced immune protection provided by ecto-CRT expression against B16.

## NCR1 recognition of ecto-CRT mediates immune protection after doxorubicin

Next, we evaluated the importance of the NCR1-ecto-CRT interaction in immune-mediated clearance of drug-treated tumors. WT and *Ncr1^−/−^* mice bearing WT B16 tumors were treated twice with doxorubicin (DOX), an ICD-inducing drug that induces ecto-CRT on B16^37^, which we verified (**Extended Data Fig. 7a**). CRT externalization induced by DOX was secondary to ER stress; it was strongly inhibited by the PERK pathway inhibitor GSK2606414, less strongly inhibited by the IRE1 inhibitor 4μ8C and not affected by the S1P inhibitor PF-429242 (**Extended Data Fig. 7b**). Although DOX significantly reduced B16 tumor growth in WT mice, its antitumor effect was not significant in *Ncr1^−/−^* mice (**Extended Data Fig. 7c**). Although there was no significant difference in the numbers of tumor-infiltrating immune cells in untreated or DOX-treated WT and *Ncr1^−/−^* mice 3 weeks after tumor implantation, increases in GzmB and PFN staining, IFN-γ and TNF-α secretion and degranulation of tumor-infiltrating NK seen in WT mice did not occur in *Ncr1^−/−^* mice (**Extended Data Fig. 7d-g**). DOX also increased CD8^+^ TIL IFNγ secretion more strongly in WT than *Ncr1^−/−^* mice. Thus, NKp46 enhances NK anti-tumor immunity after treatment with an ICD drug.

## NKp46 recognizes some senescent cells via ecto-CRT

NK recognition of senescent cells contributes to anti-tumor immunity^38^. However, how NK recognize senescent cells is unknown. To investigate which NK activating receptors recognize senescent cells, senescence was induced in the KRAS mutant human lung cancer cell line A549 by treatment with the MEK inhibitor trametinib and the CDK4/6 inhibitor palbociclib (T+P), which induces senescence and NK susceptibility^38^. ER stress is known to regulate cellular senescence in RAS-driven tumors^39^. As expected, T+P upregulated senescence-associated beta-galactosidase (SA-β-gal) activity in A549 (**Extended Data Fig. 8a**) and also induced ER stress, assessed by measuring *XBP1* splicing and *BIP* and *CHOP* mRNAs (**Extended Data Fig. 8b**). T+P-treatment also strongly induced surface expression of ecto-CRT, the NKG2D ligands (MICA/B) and intercellular adhesion molecule–1 (ICAM-1)^38^ (**Extended Data Fig. 8c**). High surface expression of ecto-CRT and MICA/B suggested that NK killing of senescent A549 might be mediated by both NKp46 and NKG2D. We therefore measured NK killing of senescent A549 by the human NK line YT in the presence of blocking antibodies to NK activating receptors, NKp46, NKG2D, or NKp30, or CRT or control antibody (**Extended Data Fig. 8d**). In the presence of control antibody, T+P potently induced A549 killing. Blocking antibodies to NKp46, CRT or NKG2D significantly reduced NK killing, but anti-NKp30 had no significant effect. Blocking both NKp46 and NKG2D reduced NK killing to background levels, indicating that NKp46 and NKG2D are the main receptors involved in NK killing of T+P-treated senescent A549. To confirm the role of NKp46 in killing senescent cells, killing by YT that express endogenous *NCR1* or were knocked out for *NCR1* was compared (**Extended Data Fig. 8e**). There was little killing of untreated A549 by *NCR1* deficient or sufficient YT. *NCR1* sufficient YT potently killed T+P-treated A549 and killing was significantly reduced by anti-CRT. By contrast *NCR1* deficient YT were much less potent at eliminating T+P-treated A549. Moreover, anti-CRT did not affect *NCR1^−/−^* YT killing of senescent A549. Thus, T+P-treated senescent A549 externalize CRT, which activates NKp46 on human NK and NKp46-dependent killing that depends on ecto-CRT.

To determine whether murine NCR1 similarly recognize ecto-CRT on T+P-treated senescent mouse lung cancer cells, murine splenic NK were used to kill KP lung cancer cells, isolated from p53 deficient mice expressing mutant KRAS^40^. As for human lung cancer cells, T+P treatment induced senescence in KP as measured by SA-β-gal (**Fig. 4e**), ER stress (**Fig. f**) and expression of ecto-CRT and surface RAE1, a mouse ligand of NKG2D (**Fig. 4g**). Splenic NK from WT mice barely killed untreated KP but were highly cytolytic against T+P-treated KP (**Fig. 4h**). However, splenic NK from *Ncr1^−/−^* and *Klrk1^−/−^* mice deficient in NCR1 and NKG2D, respectively, had impaired killing compared to NK proficient in both NK activating receptors, indicating that both NK activating receptors recognized T+P-treated KP. T+P strongly suppressed KP tumors in WT mice but tumor growth was not significantly slowed in *Ncr1^−/−^* mice (**Fig. 4i**). NCR1 had a more potent role in controlling T+P-treated KP tumor growth than NKG2D as T+P-treated KP tumors still grew significantly more slowly in *Klrk1^−/−^* mice (*P*=0.04). Thus, T+P-treated senescent human and mouse RAS-driven lung tumors are recognized as stressed cells by NKp46/NCR1 through ecto-CRT and by NKG2D. Although the numbers of tumor-infiltrating immune cells did not significantly change in T+P-treated mice (**Extended Data Fig. 9a**), tumor infiltrating NK in T+P-treated WT mice had significantly more PFN and TNF-α staining, but these differences were no longer significant in *Ncr1^−/−^* mice and only TNF-α staining increased significantly in *Klrk1^−/−^* mice (**Fig. 4j**). These functional parameters on CD8^+^ TIL did not significantly change after T+P treatment (**Extended Data Fig. 9b,c)**. Thus, NKp46 and NKG2D recognition of T+P-treated senescent lung cancers enhances NK anti-tumor functions and tumor control, but NCR1 may be somewhat more important.

To determine whether NKp46 recognition of senescent cells depends on ER stress and ecto-CRT, KP were treated with two other inducers of senescence - CuSO_4_ and 4-phenyl-3-butenoic acid (4-PBA). Treatment with CuSO_4_, but not 4-PBA, is reported to induce ER stress^41,42^. As expected, both compounds increased senescence assessed by SA-β-gal activity (**Extended Data Fig. 10a**), but only CuSO_4_ induced ER stress assessed by increased *Xbp1* splicing and *Bip* and *Chop* mRNA (**Extended Data Fig. 10b**). Ecto-CRT and surface RAE1 were both strongly induced by CuSO_4_, but weakly, but significantly, induced by 4-PBA (**Extended Data Fig. 10c**). CuSO_4_ significantly increased in vitro KP killing by splenic NK isolated from WT mice and killing of CuSO_4_-treated KP was significantly reduced by splenic NK from both *Ncr1^−/−^* and *Klrk1^−/−^* mice, indicating that both activating receptors contributed to killing (**Extended Data Fig. 10d**). 4-PBA treated KP cells were weakly killed by splenic NK and this weak killing depended only on NKG2D since T+P treatment did not increase killing by *Klrk1^−/−^* NK but did increase killing by *Ncr1^−/−^* NK. Thus, NK do not efficiently recognize and kill all senescent cells; NK killing of senescent tumor cells is enhanced by ER stress and NKp46/NCR1 recognition of ecto-CRT in the tumor.

## Discussion

Here we show that NKp46/NCR1 directly recognizes ecto-CRT on the surface of cells undergoing ER stress triggered by multiple inducers (ZIKV, Tu, ICD-inducing chemotherapy (OP, DOX), senescence). The identification of ER stress and its cell surface indicator ecto-CRT as the endogenous ligand of the evolutionarily most ancient and conserved NK activating ligand NKp46 makes sense and fits the hypothesis that NK activating receptors sense cellular stress^43^. NKG2D, which recognizes other cellular stresses^44,45^, also contributed to NK activation by senescent cells and other chemotherapy drugs (CP). The identification of ecto-CRT as the endogenous ligand of NKp46 by pulldown and mass spectrometry was confirmed by in vitro binding experiments using recombinant proteins and by cellular experiments using CRT blocking antibody, *CALR* knockout and knockdown, ectopic expression of GPI-CRT on target cells and by knocking out *NCR1/Ncr1* in a human NK line and in mice. Ectopic ecto-CRT expression or induction of ecto-CRT by DOX treatment of a melanoma cell line or T+P treatment of a RAS-driven lung cancer line greatly reduced tumor growth in an *Ncr1*-dependent manner. Although the numbers of tumor-infiltrating immune cells were not significantly increased in WT vs *Ncr1^−/−^* mice, the presence of the NKp46 activating receptor and its ligand on the tumor increased NK TIL anti-tumor functionality (expression of cytotoxic granule proteins, degranulation and cytokine secretion). Although NK have been shown to play an important role in recruiting DC into tumors, which enhances anti-tumor immunity^46^, in this study *Ncr1* sufficiency did not appreciably change DC infiltration or tumor phagocytosis.

ER stress is a feature of many disease processes^47^. Infection with many viruses, including both DNA and RNA viruses, induces ER stress and some viruses have developed strategies to counter this host response. It is likely that cells infected with other ER-stress-inducing viruses, including other flaviviruses besides ZIKV, like dengue virus and hepatitis C, which replicate in the ER, will expose ecto-CRT and be recognized by NKp46. In fact high expression of NKp46 on liver NK has been linked to better outcome in hepatitis C infection^48^. Stresses in tumors, including hypoxia and disruption of proteostasis in lymphomas and myelomas, also cause ER stress, which could recruit and activate NK immune surveillance. Metabolic dysregulation in diabetes and obesity also causes ER stress^49^. Future studies of cell surface CRT and its recognition by NKp46 on NK and ILC in different disease settings will enhance understanding of the contribution of this innate immune response to disease control and pathogenesis. Our experiments inhibiting different ER stress pathways in ZIKV-infected JEG3 and DOX-treated B16 did not identify a shared required ER stress pathway for CRT exposure, suggesting that additional work is needed to understand the mechanistic basis of CRT exposure during ER stress.

Irradiation and some anticancer drugs, such as OP and anthracyclines, like DOX, induce ICD, which plays a critical role in whether cancer treatments are curative (at least in mice) by eliminating cancer cells not directly killed by the treatment^50^. Tumor ER stress and ecto-CRT are the hallmarks of ICD, although other factors including secretion of alarmins, such as ATP, HMGB1 and annexin A1, have also been implicated. The dominant immune protective mechanism in ICD has been thought to be ecto-CRT acting as an “eat me” signal that promotes tumor cell phagocytosis by dendritic cells, which in turn activate anti-tumor CD8^+^ T cells by cross-priming to kill tumor cells that survive chemotherapy^51^. Our findings here suggest that NKp46-mediated NK killing of ecto-CRT-expressing cancer cells is also important in tumor immune surveillance and ICD-related tumor immune control. In fact, in some situations, NK recognition of ecto-CRT may be the most important immune mechanism responsible for ICD protection. For example, protection afforded by ectopic expression of GPI-CRT or DOX treatment in B16 melanoma-bearing mice or by T+P treatment of KP lung cancer-bearing mice was largely abrogated in *Ncr1* deficient mice, suggesting that NKp46-mediated NK recognition and killing drive immune protection in those models.

NK play an important role in tumor immune surveillance^52^. However, their role in tumor immune defense has been overshadowed by a focus on anti-tumor CD8^+^ T cells. Like T cells, NK immunity is compromised as tumors develop - generally a relatively small number of NK are found infiltrating tumors and NK also become “exhausted” as tumors progress^53^. Nonetheless, harnessing NK immunity for cancer immunotherapy using adoptively transferred NK or CAR-NK has recently shown promise, especially for hematopoietic cancers^54^. The limited understanding of the endogenous signals that activate NK has impeded research and translational efforts to augment NK immunity. Our identification here of ecto-CRT as an important trigger for NK recognition of cancer cells, undergoing ER stress, senescence or after chemotherapy or irradiation, should open up new approaches for exploiting and enhancing NK immune defenses against viral infection and cancer. Developing therapeutic strategies to enhance NK tumor immunity will be facilitated by a better understanding of the molecular mechanisms responsible for ecto-CRT exposure.

## EXTENDED DATA FIGURES

**Extended Data Fig. 1.**
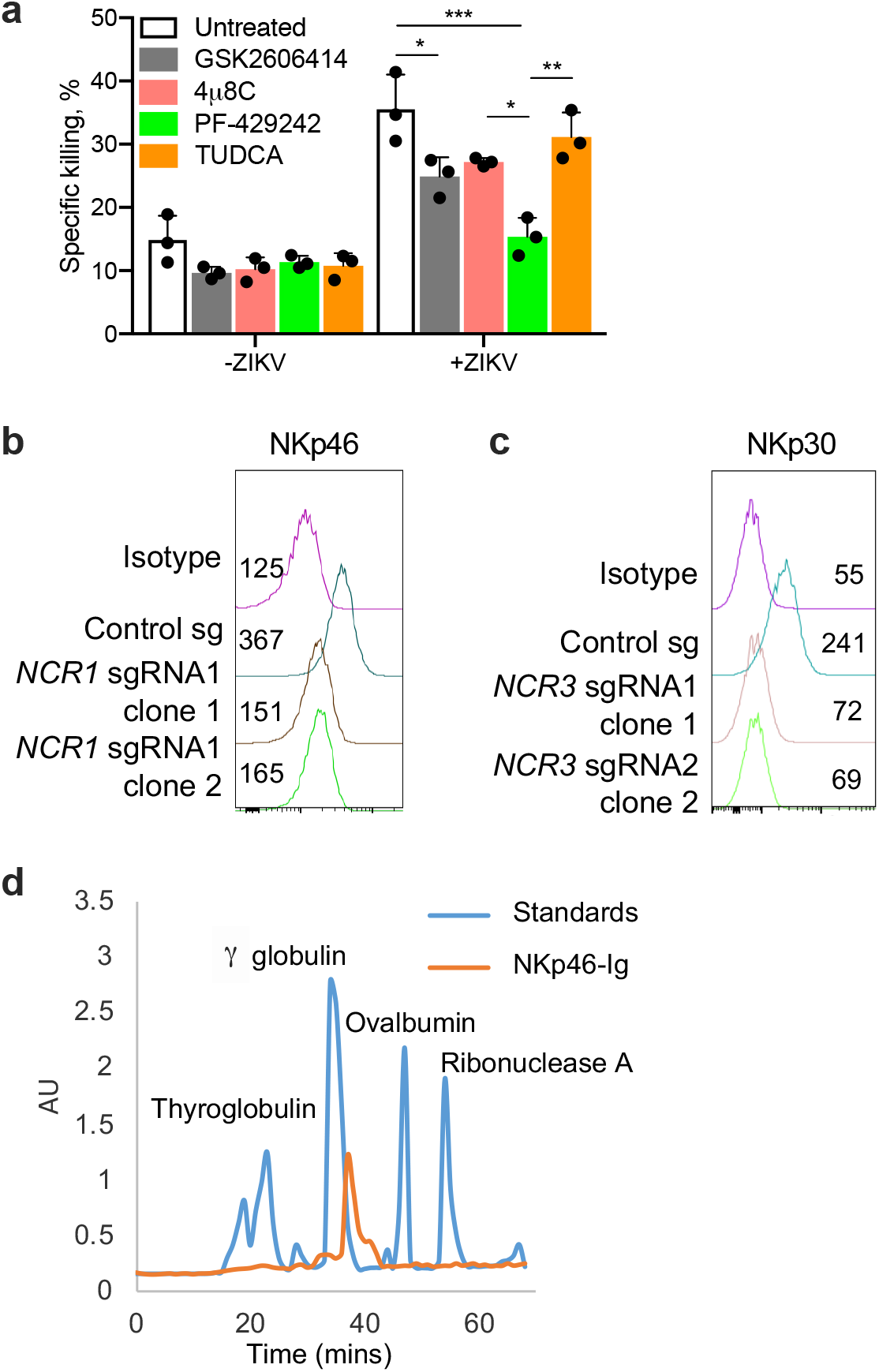
**a,** Effect of ER stress inhibitors (GSK2606414, 4u8C, PF-429242 and TUDCA) on human peripheral blood NK killing of ZIKV-infected JEG-3 assessed by 8 h ^51^Cr release assay using an E:T ratio of 10:1 (n=3). **b,c** Representative flow cytometry histograms of NKp30 (b) and NKp46 (c) surface expression on WT or CRISPR knockout clones of human YT NK cells. MFI is indicated. **d,** Gel filtration analysis of NKp46-Ig and protein standards separated on a Superdex 2000 Tricorn 10/600 column showing NKp46 migrates predominantly as a dimer and is not aggregated. Data in (a) are mean ± SEM of three technical replicates. Statistics by one-way ANOVA (**a**) *P*: *<0.05, **<0.01, ***<0.001.

**Extended Data Fig. 2.**
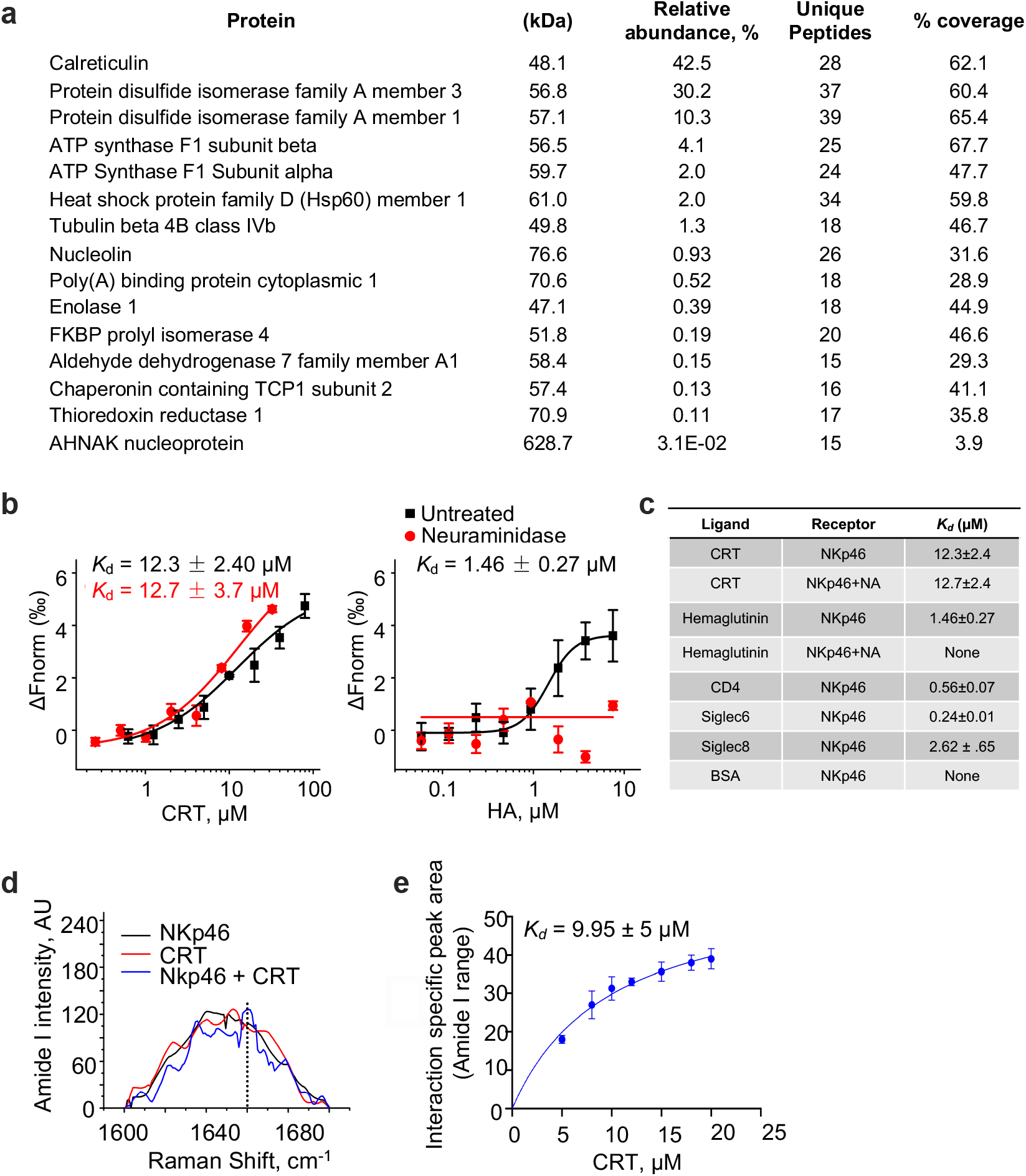
NKp46 binds to calreticulin. **a,** Proteins identified by mass spectrometry analysis of the cross-linked high molecular band in the NKp46-Ig pulldown of the membrane fraction of ZIKV-infected JEG-3, listed in order of ion abundance. **b,** Binding of neuraminidase treated (red) or untreated (black) Alexa647-labeled NKp46-Fc to His-tagged CRT (left) or His-tagged hemagglutinin (HA) (right) analyzed by microscale thermophoresis (MST, dissociation constants K_d_ are indicated). **c**, Binding Kds measured by MST of CRT with NKp46 and other reported ligands. Data are mean±s.d. of three technical replicates. **d,** Raman spectra of normalized intensities (arbitrary units, a.u.) for recombinant NKp46 and CRT in the amide I range (1600-1700 cm^−1^) recorded individually and then after mixing. The spectrum of the mixture shows a new spectral feature (peak at, dotted line) potentially indicating NKp46-CRT binding (left). Change in the peak area at 1658 cm^−1^ plotted against increasing CRT concentrations. The peak areas were obtained by deconvolution of the spectral read-outs (n=3) and the plotted data were subjected to hyperbolic fitting to derive the NKp46-CRT dissociation constant.

**Extended Data Fig. 3.**
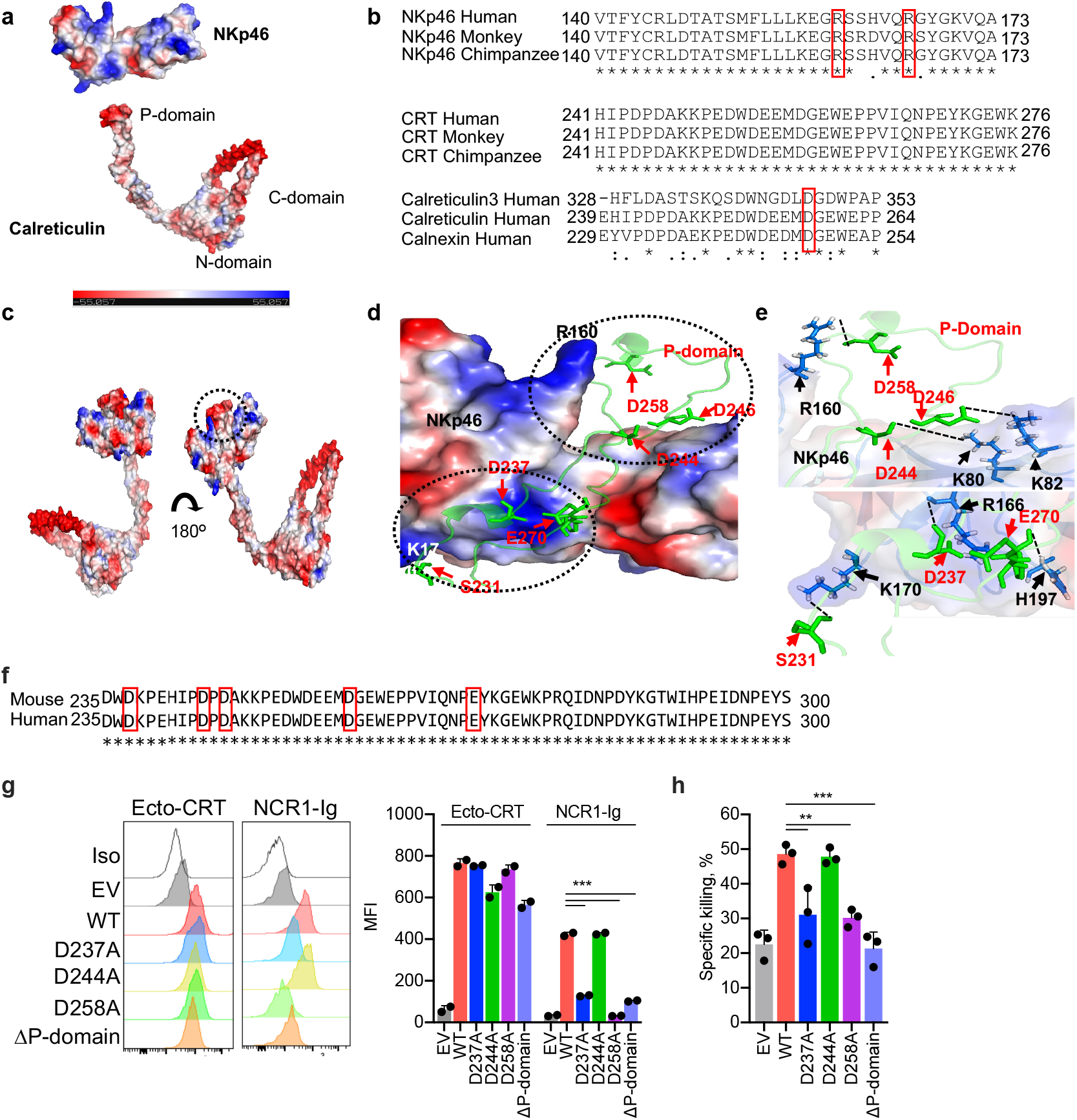
Modeling of the interaction of the calreticulin P-domain with NKp46. **a,** Molecular surface representation of NKp46 (accession no. O76036) and CRT (accession no. P27797) based on sequence derived 3D structure generated by I-TASSER program **b,** Sequence alignment of human NKp46 (top) and CRT (middle) proteins (accession nos. O76036, P27797), monkey (Q8MIZ9, Q4R6K8) and chimpanzee (Q08I01, H2QFH8), respectively. Conserved residues important in binding are labeled with a red box. Sequence alignment of the tip module of P domains of the CRT/CNX family (bottom). The Asp258 residue required for binding is labeled with a red box. **c,** Docked complex of NKp46 and CRT represented as a surface assembly; surface charge is shown for the entire complex (scale bar representing the charge gradient from negative (red) to positive surface charge (blue)). Circled region represents the binding pocket, which is further magnified in (**d**). **d,** Magnified interaction of NKp46 (surface charge representation) and CRT P-domain (in stick and ribbon representation) showing specific Asp residues of the P-domain in the positively charged cleft of NKp46. NKp46 R160 is marked in the positively charged region of the NKp46 cleft. **e,** Magnification of the circled regions at the top and bottom of the binding cleft in **(d)** showing salt bridges and hydrogen bonds (dashed black lines) between NKp46 residues (blue sticks and black labels) and CRT (green sticks and red labels). **f,** Sequence alignment of the P domains of mouse and human CRT. The Asp and Glu residues implicated in binding are labeled with a red box. **g,** Representative flow cytometry histograms of CRT surface expression and NCR1-Ig binding (left) and mean fluorescence intensity (MFI, right) of 2 samples of B16 cells stably transfected with empty vector (EV) or WT or mutated GPI-CRT. Untransfected B16 display little ecto-CRT. **h,** Killing by splenic NK from WT mice (n=3) of B16 stably transfected with empty vector (EV) or WT or mutated GPI-CRT (8 h ^51^Cr release assay, E:T ratio 10:1). Data shown are representative of 3 experiments. Statistics by one-way ANOVA (g,h). *P*: **<0.01; ***<0.001.

**Extended Data Fig. 4.**
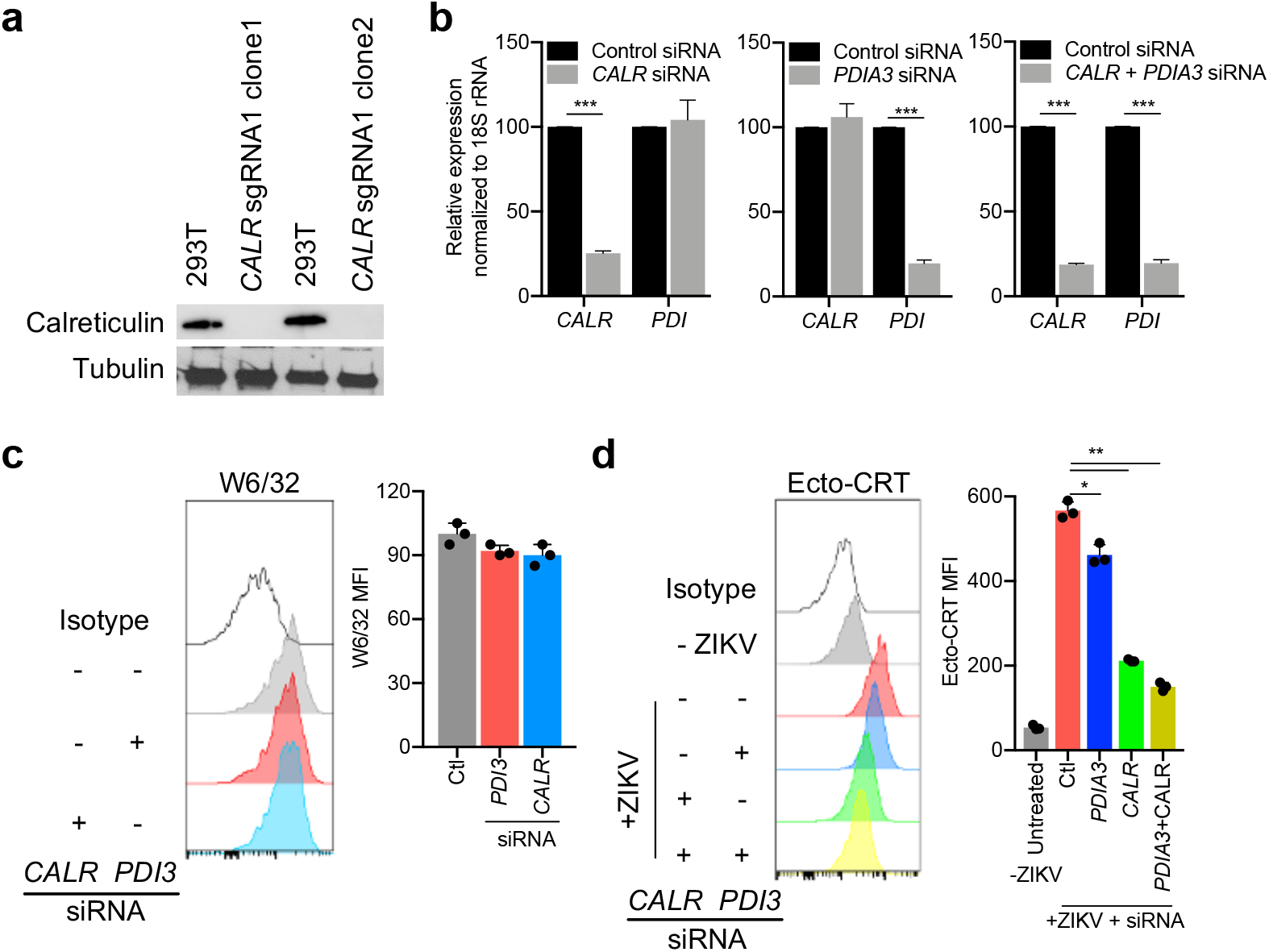
Knockout and knockdown of *CALR* and *PDI3A*. **a,** Immunoblot of CRT in WT or CRISPR knockout HEK293T clones compared to tubulin loading control. **b,** qRT-PCR analysis of *CALR* and *PDI* expression after siRNA-mediated knockdown of indicated genes in JEG-3, normalized to *18S rRNA*. **c,** Representative flow cytometry histograms of MHC class I surface expression analyzed by W6/32 antibody staining on JEG-3, knocked down for *CALR, PDI3 or* with nontargeting (Ctl) siRNAs. MFI of 3 samples is indicated (right). **d,** JEG-3, knocked down for *CALR* and/or *PDI3 or* with nontargeting (Ctl) siRNAs were infected or not with ZIKV and analyzed for ecto-CRT by flow cytometry. Representative histograms are shown at left and MFI of 3 samples is indicated at right. Shown are representative of 3 experiments. Statistics by non-parametric unpaired *t*-test (b) or one-way ANOVA (c,d). *P*: *<0.05; **<0.01; ***<0.001.

**Extended Data Fig. 5.**
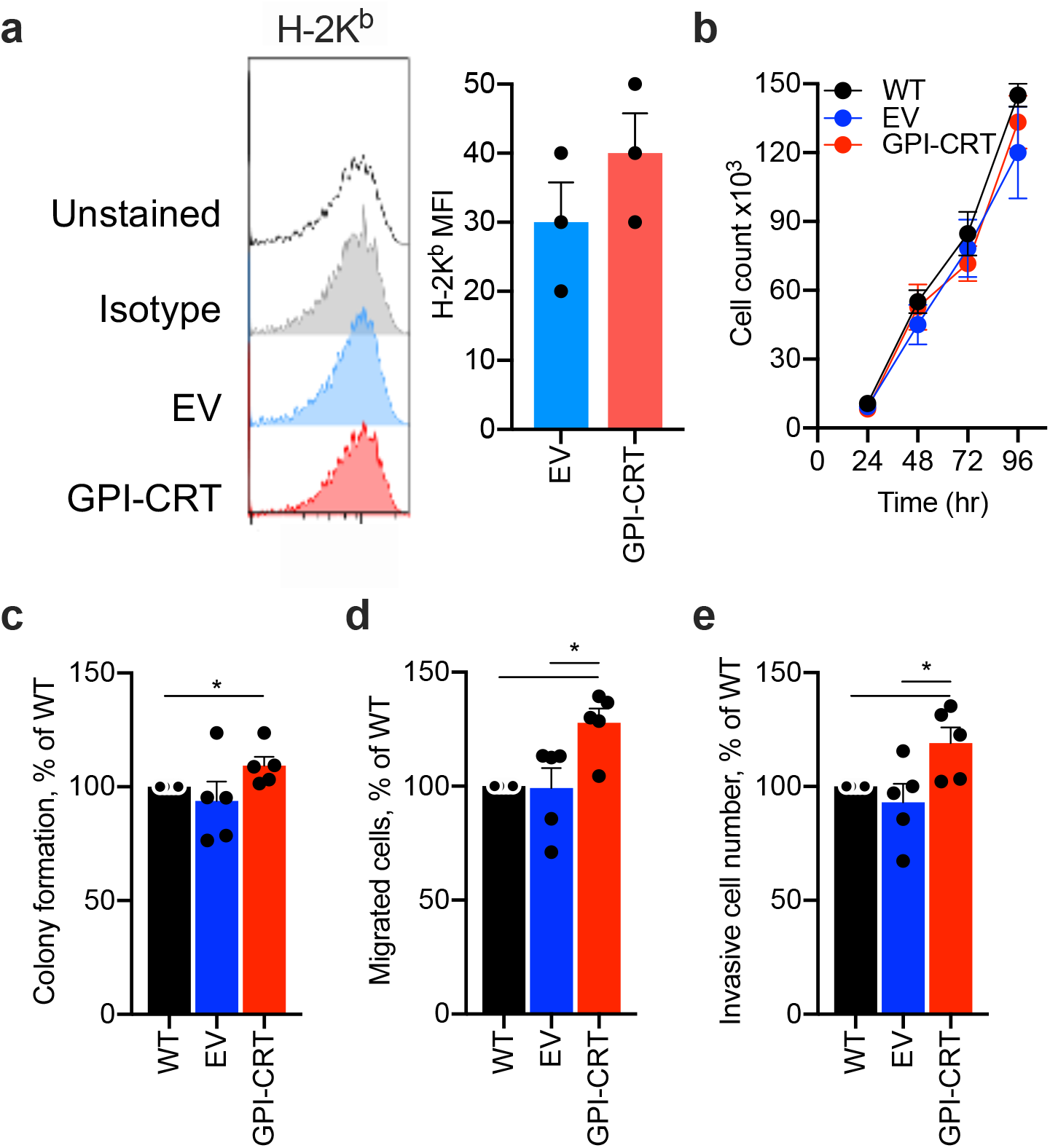
Ectopic expression of GPI-linked CRT in B16 does not alter in vitro cell proliferation but increases colony formation, invasivity and migration across a membrane. B16 were not transfected or were stably transfected with empty vector (EV) or with an expression plasmid for GPI-CRT and analyzed for H-2K^b^ expression by flow cytometry (n=3) (**a**), cell proliferation (**b**), colony formation (**c**), invasion through a Transwell (**d**) and migration through a Transwell membrane in response to serum (**e**) (b-e, n=5). **a,** Representative flow cytometry histograms of H-2K^b^ surface expression on B16 (left); graph at right shows mean±SEM MFI of multiple samples. **c,** For colony formation assay, 500 cells were seeded in 10 mm^2^ plates and stained with crystal violet after 10 days. **d,** Cell invasion was assessed by plating 1×10^5^ cells in Matrigel on the upper chamber membrane and measuring cell counts in the bottom chamber surface after 24 h. **e,** Migration was assessed by seeding cells in the top chamber of a Transwell in serum-free medium and counting the cells that had migrated 24 h later to the bottom chamber, which contained 10% FCS. Shown are mean ± SEM of at least 3 replicates. Statistics by non-parametric unpaired *t*-test (a), area under the curve, followed by one-way ANOVA (b) and one-way ANOVA (c-e). *P*: *<0.05.

**Extended Data Fig. 6.**
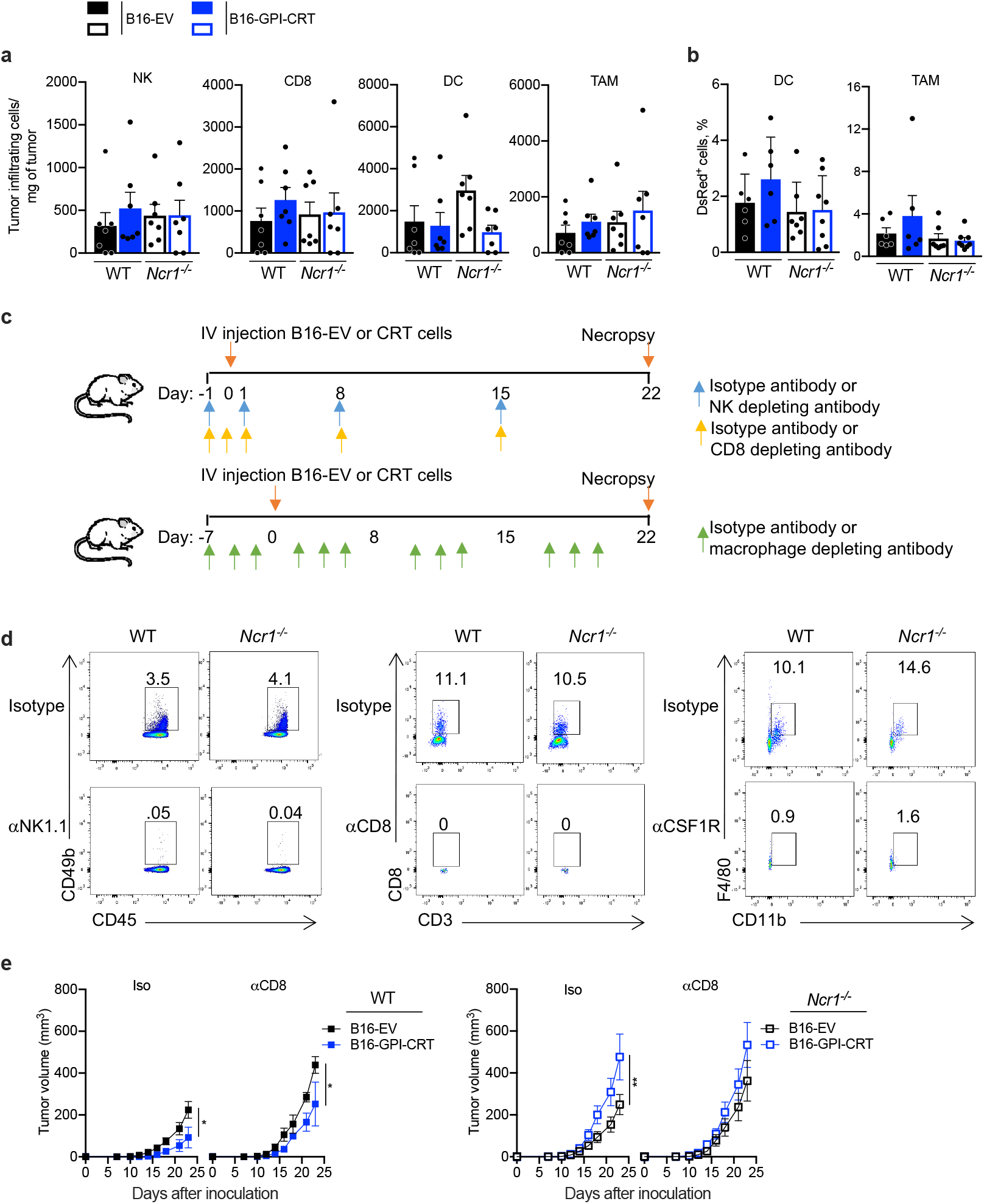
Ectopic expression of ecto-CRT in B16 does not significantly change numbers of tumor-infiltrating cells but suppresses tumor growth in an NCR1-dependent, but CD8^+^ T cell-independent, manner. Empty vector (EV) or GPI-CRT expressing B16 tumors were implanted sc in WT and *Ncr1^−/−^* mice (n=7/group) and mice were sacrificed 24 d later (extended data for Fig. 4-c). **a,** Numbers of tumor infiltrating cells. **b,** Percentage of DsRed^+^ DC and TAM infiltrating DsRed^+^ B16 tumors **c,** Schematic of metastasis experiment in Fig. 4d. **d,** Representative flow cytometry plots of mouse blood mononuclear cells, obtained on day 3 after tumor implantation, showing NK (left), CD8^+^ T (middle) and monocyte (right) depletion with cell type-specific antibody compared to isotype control antibody. The integrin CD49b is a pan-NK marker. **e,** Tumor growth in WT (left) or *Ncr1^−/−^* (right) mice injected sc with empty vector (EV) or GPI-CRT-overexpressing B16 that were treated with an isotype control antibody or depleted of CD8 T cells prior to tumor implantation and during the course of the experiment (n=5/group). Graphs in (**a,b**) show mean ± SEM of 3 technical replicates and at least 5 biological samples representative of 2 independent experiments; graphs in (**e**) are mean ± SEM. Statistics calculated by non-parametric one-way ANOVA followed by Tukey’s post-hoc test for areas under curves (e) and unpaired Student’s t-test (a,b). *P*: *<0.05; **<0.01.

**Extended Data Fig. 7.**
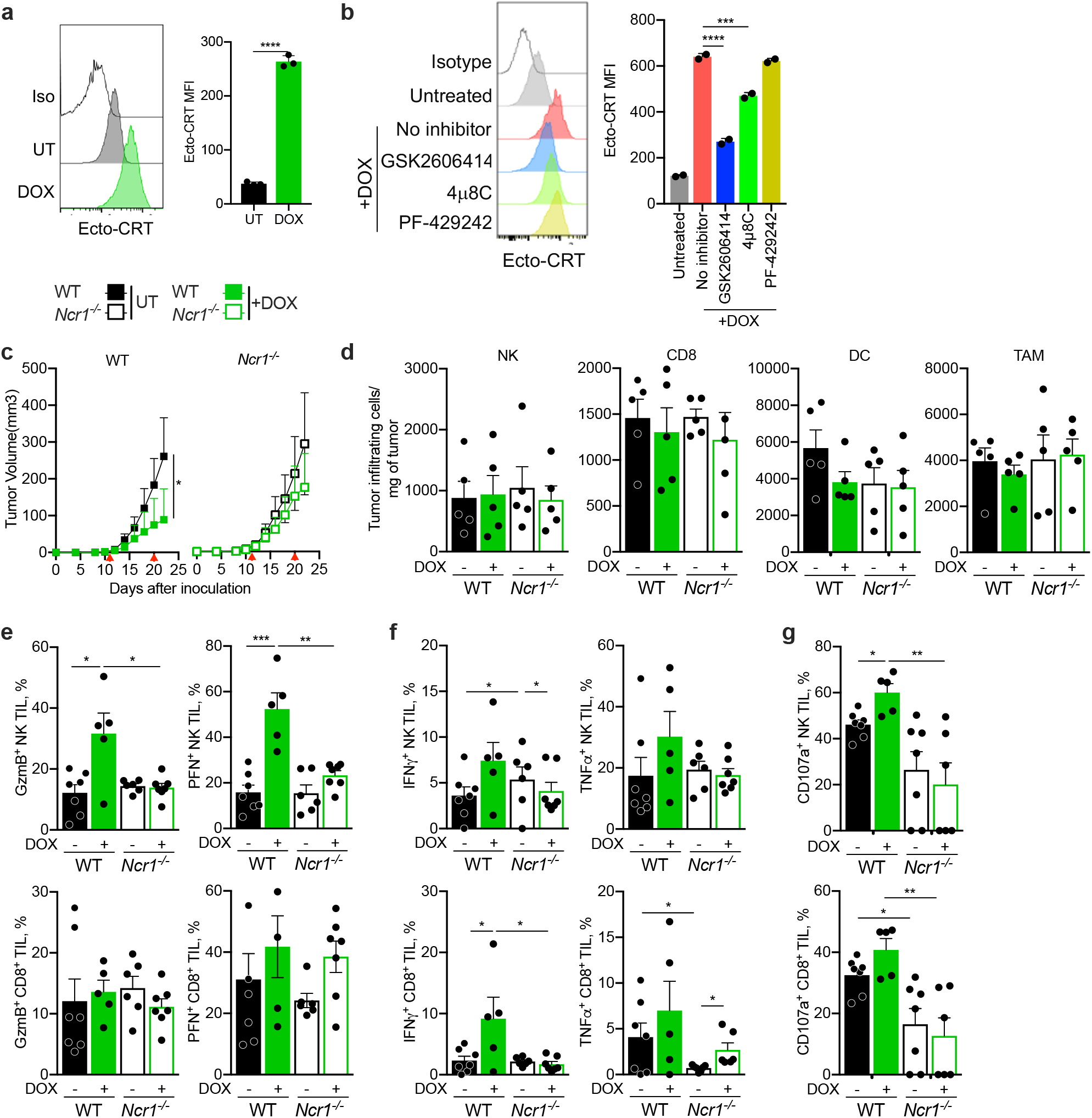
NCR1 enhances doxorubin suppression of B16 tumors. **a,** Representative flow cytometry histograms of ecto-CRT (left) and mean±SEM ecto-CRT mean fluorescence intensity (MFI, right) of B16 treated in vitro for 24 h with doxorubicin (DOX) or left untreated (UT). **b,** Representative flow cytometry histograms of ecto-CRT expression on UT and DOX-treated B16 that were pretreated or not with indicated ER stress inhibitors. **c-g,** B16 cells were injected sc into WT or *Ncr1^−/−^* mice (n = 7/group). and animals were treated with DOX iv 12 and 20 d after tumor implantation (red arrows). Tumor growth (**c**), numbers of tumor-infiltrating immune cells **(d)** and functional markers of NK (top) and CD8^+^ (bottom) TIL were assessed at time of sacrifice (cytotoxic granule protein expression (**e),** cytokine secretion **(f)** and degranulation **(g)** after PMA + ionomycin stimulation ex vivo). Data in bar graphs show mean ± SEM of at least 5 biological samples and are representative of 2 independent experiments. Statistics calculated by non-parametric one-way ANOVA followed by Tukey’s post-hoc test for areas under curves (c) and two-tailed Student’s t-test (a,b,d,e,f). *P*: *<0.05; **<0.01; ***<0.001, **** <0.0001.

**Extended Data Fig. 8.**
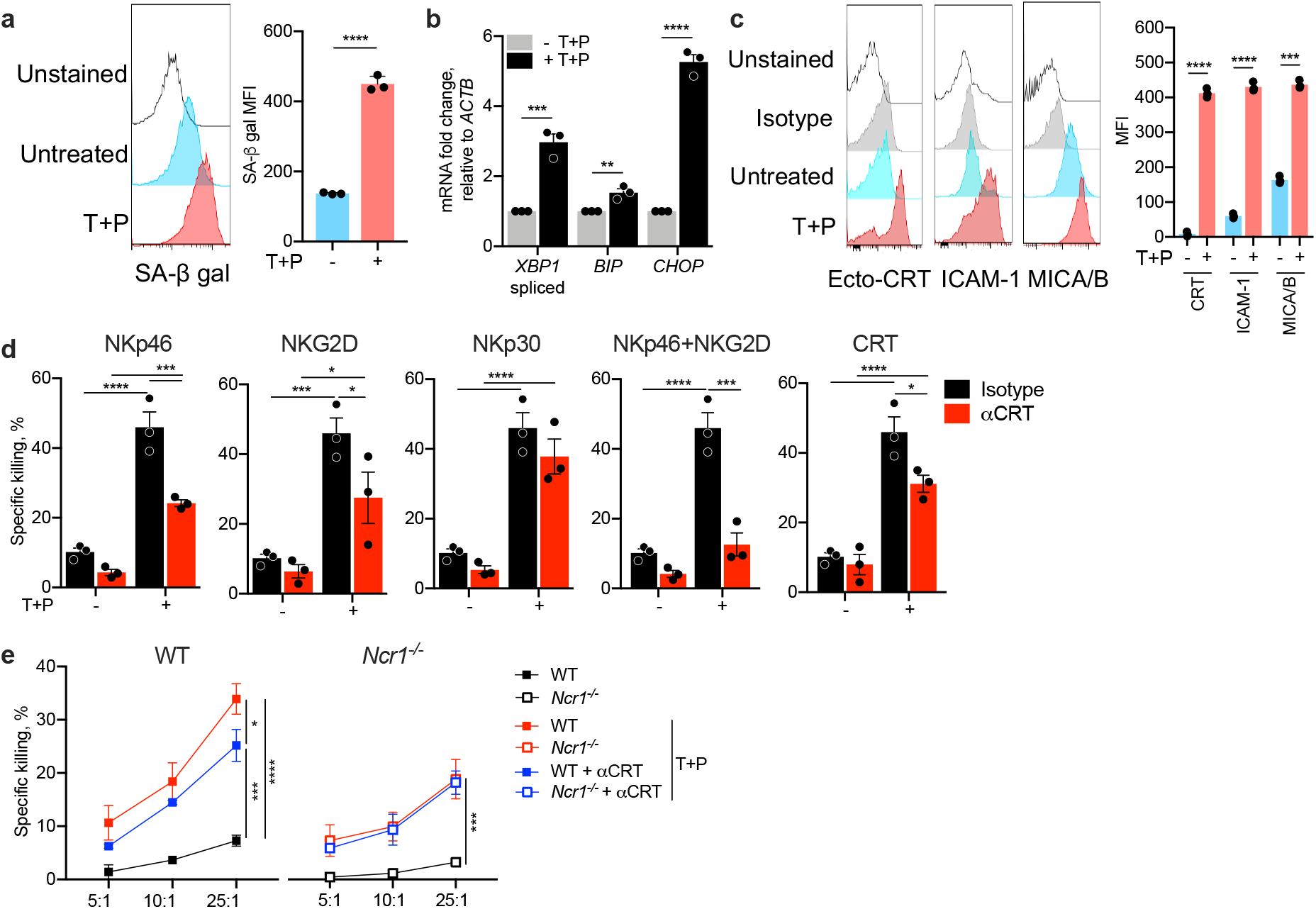
Trametinib and palbociclib induces senescence in human A549 lung cancer cells and activates NK through NKp46 and NKG2D. **a,** Representative flow cytometry histograms of β-galactosidase activity (SA-β-gal) in A549 that were untreated or treated with trametinib and palbociclib (T+P) (left) and mean±SEM mean fluorescence intensity (MFI, right) of multiple samples (n=3). **b,** ER stress, assessed by qRT-PCR assay of *XBP1* splicing and *BIP* and *CHOP* mRNA, in untreated and T+P-treated A549 (n=3). **c,** Representative flow cytometry histograms of CRT, ICAM1 and MICA/B expression on untreated and T+P-treated A549 (left) and mean±SEM MFI (right) (n=2). **d,** Effect of NK receptor blocking antibodies (NKp46, NKG2D and NKp30) or anti-CRT on YT killing of untreated or T+P-treated A549 (8 h ^51^Cr release, E:T ratio 25:1, n=3). **e,** Specific killing of untreated or T+P-treated A549 by YT, knocked out or not for *NCR1* in the presence or absence of CRT blocking Ab (8 h ^51^Cr release, E:T ratio 25:1). Bar graphs in (a-d) show mean ± SEM of at least 3 independent experiments and statistics by unpaired one-way ANOVA. Statistics were calculated by non-parametric one-way ANOVA followed by Tukey’s post-hoc test for area under curves (e). *P*: *<0.05; **<0.01; ***<0.001, **** <0.0001.

**Extended Data Fig. 9.**
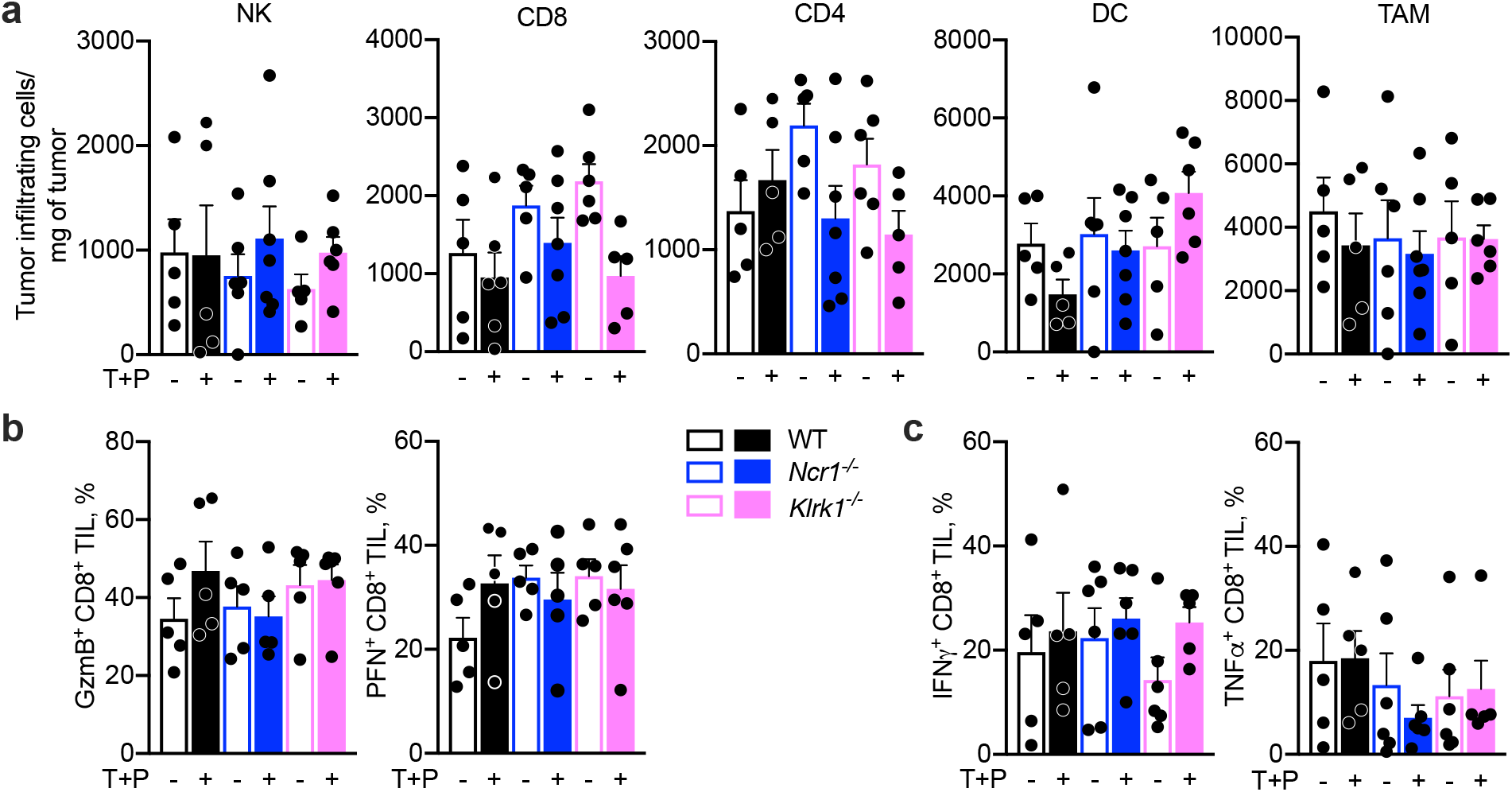
Trametinib and palbociclib treatment of WT and NRC1 or NKG2D deficient mice bearing subcutaneous KP tumors does not affect the numbers of tumor-infiltrating immune cells or the functional phenotype of tumor-infiltrating CD8^+^ T cells. KP cells were injected sc into WT, *Ncr1^−/−^* or *Klrk1^−/−^* mice (n = 5-7/group) and animals were treated with T+P by oral gavage 13-18 d after tumor implantation. (Supplemental data for Fig. 4i-k.) Tumor-infiltrating cells were analyzed at the time of sacrifice (23 d post tumor implantation). **a,** Number of tumor-infiltrating immune cells. **b,c,** CD8^+^ TIL expression of cytotoxic granule proteins (**b)** and cytokine production (**c**). Bar graphs show mean ± SEM of at least 3 independent experiments and statistics by unpaired one-way ANOVA. There were no statistically significant differences between the groups.

**Extended Data Fig. 10.**
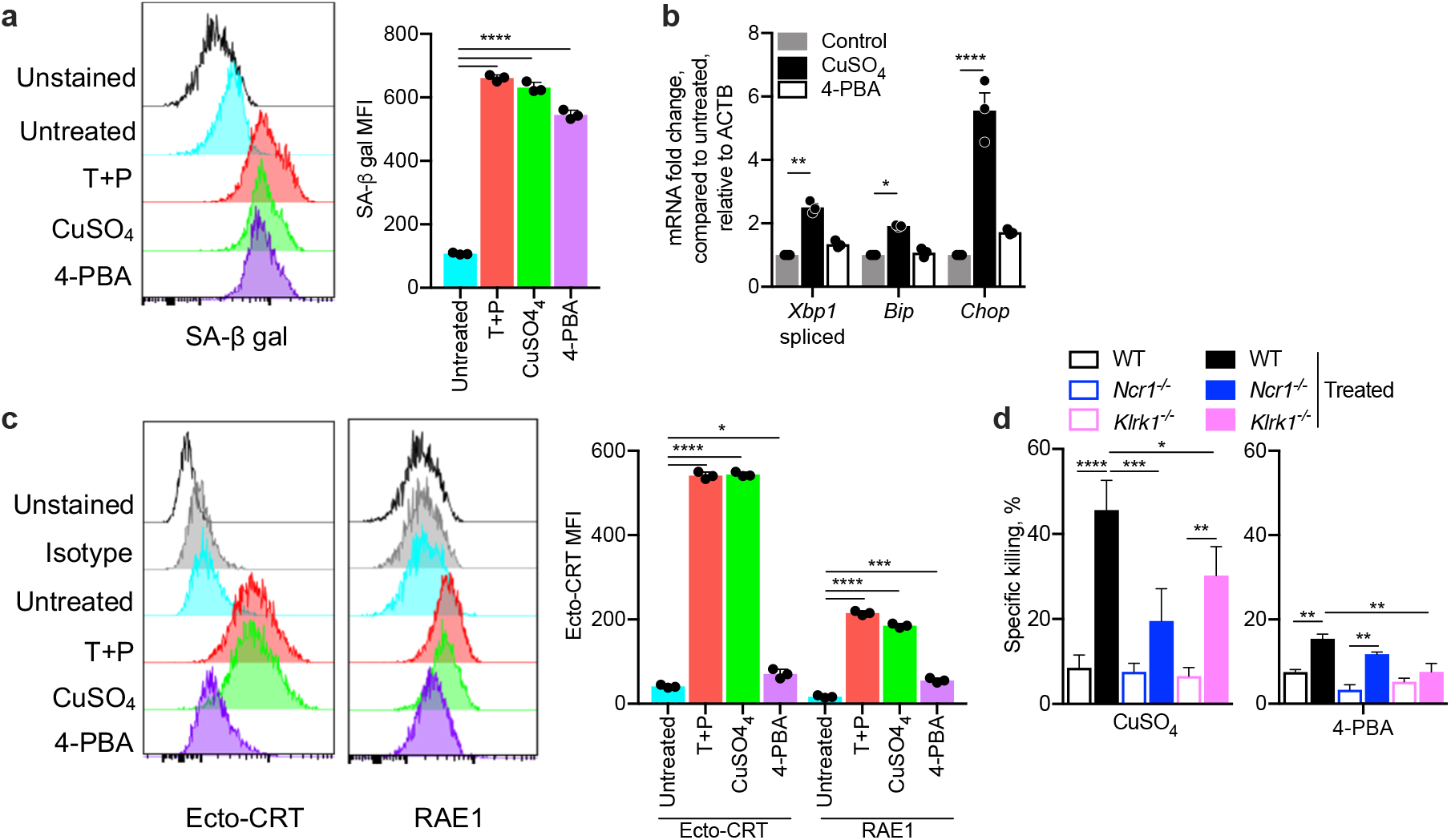
Senescence inducer CuSO_4_, which causes ER stress, activates NCR1- and NKG2D-dependent NK killing but 4-PBA, which induces senescence without ER stress, does not. **a,** Representative flow cytometry histograms of β-galactosidase activity (SA-β-gal) (left) and mean±SEM mean fluorescence intensity (MFI, right) of multiple samples (n=3) of KP cells that were untreated or treated with CuSO_4_ or 4-PBA. **b,** ER stress, assessed by qRT-PCR assay of *Xbp1* splicing and *Bip* and *Chop* mRNA, in untreated and CuSO_4_ or 4-PBA treated KP (n=3). **c,** Representative flow cytometry histograms of CRT and RAE1 expression on KP untreated and treated with CuSO_4_ or 4-PBA (left); mean±SEM MFI of multiple samples (right, n=3). **d,** Killing of KP cells that were untreated or treated with CuSO_4_ or 4-PBA by splenic NK from WT, *Ncr1^−/−^* or *Klrk1^−/−^* mice (4 h ^51^Cr release, E:T ratio 20:1). Bar graphs show mean ± SEM of at least 3 independent experiments. Statistics by unpaired one-way ANOVA. *P*: *<0.05; **<0.01; ***<0.001, **** <0.0001.

## METHODS

### Cell lines

JEG-3, A549, 721.221, HFF, Vero, HEK293T and B16 were from ATCC. YT was a kind gift of Z. Brahmi, Indiana University. The KP cell line was obtained from T. Jacks (Koch Institute, MIT)^40^. Cell lines were recent passages and were periodically tested for mycoplasma contamination; flow cytometry was used to confirm cellular identity by cell surface markers. 721.221 and YT were cultured in RPMI 1640 and JEG-3, A549, HFF, Vero, 293T, B16, and KP were cultured in DMEM. All media were supplemented with 10% FCS, 1% Pen/Strep and 1% L-glutamine. Stable ecto-CRT expressing B16 were generated as described in ^34^. All stably transfected clones were verified by surface CRT staining and flow cytometry.

### Viruses

ZIKV PRVABC59 strain (Puerto Rico, 2015) was obtained from the Arbovirus Branch of the Centers for Disease Control and Prevention (CDC). HSV-2 was obtained from ATCC. HCMV AD169 (IE-1-GFP) was a gift of Donald Coen, Harvard Medical School. ZIKV stocks were propagated in DMEM supplemented with 2% heat-inactivated FBS in Vero. ZIKV titers were determined by plaque assay on Vero incubated for 3 d using a 1.2% Avicel (RC-59INF, FMC Corporation) overlay. HCMV was propagated in HFF in DMEM supplemented with 3% FBS and supernatants were collected 8 d post infection. HCMV titers were determined by counting GFP^+^ foci on HFF 2 d post infection using a 1.2% methylcellulose (Sigma) overlay. HSV-2 was grown in Vero and viral titers were determined by plaque assay in Vero.

### Antibodies and Reagents

Antibodies and reagents used were: donkey-anti-mouse Alexa Fluor 488, LIVE/DEAD Fixable Violet Dead Cell Stain, purified mouse anti-human HLA-A, B, C (W6/32), Alexa Fluor 647 anti-human CD54 Antibody (HCD54), APC anti-human CD337 (NKp30) (P30-15), Alexa Fluor 488 anti-Hsp70 (W27), CD56 -PE and -Pacific Blue (HCD56), CD107a-PerCP-Cy5.5 (H4A3), purified mouse IgG1, IgG1-FITC, IgG1-PE, IgG1-APC, IgG1-PerCP-Cy5.5, IgG1-PE-Cy7, IgG1-APC-Cy7 (MOPC-21), purified mouse IgG2a, IgG2a-FITC, IgG2a-AlexaFluor 700 (MOPC-173, anti-mouse B220-PE (RA3-6B2), CD45-PE-Cy7 (30-F11) (BioLegend), MICA/B-Alexa Fluor 700 (159207) (R&D Systems); anti-mouse CD3e-PerCP (145-2C11), CD8 PerCP (53-6.7), IFN-γ-APC (XMG1.2) (BD); anti-mouse CD8-eFluor710 (53-6.7), CD62L-APC (MEL-14), CD44-PE (IM7), CD4-eFluor 450 (RM4-5), NKp46-PE (29A1.4) (eBioscience), FITC-AffiniPure goat-anti-human IgG (Jackson Immunoresearch); Alexa Fluor 647 anti-mouse RAE-1γ (CX1, BioLegend), FITC anti-mouse H-2K^b^ (AF6-88.5, BioLegend), Alexa Fluor 488 anti-phospho-Syk (pT348), Alexa Fluor 647 anti-phospho-CD3-zeta (pY142), PE anti-phosphotyrosine (BD Biosciences), (Santa Cruz Biotechnology), goat anti-mouse IgG Fab recombinant secondary antibody (Invitrogen), anti-NKp46 (9E2, Miltenyi Biotec) anti–mouse CD8 mAb, clone 2.43, rabbit anti–mouse NK1.1, anti-CSF1R, control antibody (LTF-2, BioXCell), rabbit anti-human CRT (ab2907, Abcam), mouse monoclonal anti-CRT (MAB38981, R&D Systems; FMC 75, Thermo); PDIA3 rabbit polyclonal Ab (A1085, Abclonal); Alexa Fluor 488 mouse IgG1 isotype control (clone MOPC-21); PE mouse IgG1 isotype control ( MOPC-21; Alexa Fluor 647 mouse IgG1 isotype control (MOPC-21), PerCP-Cy5.5 mouse IgG1 isotype control (MOPC-21), AlexaFluor 488 mouse IgG2a isotype control (MOPC-173), AlexaFluor 647 mouse IgG2b isotype control (BioLegend, MPC-11), Pacific Blue mouse IgG2b isotype control (BioLegend, MPC-11). NKp44-Ig (2249-NK), NKG2D-Ig (1299-NK), NKp46-Ig (1850-NK), NCR1-Ig (2225-NK) fusion proteins were from R&D Systems; NKp30-Ig (Acrobiosystems), hemagglutinin His-tag (SinoBiological), human CD4-His tag, Siglec8-His tag (Acrobiosystems), Siglec6-His tag (Creative Biolabs), 4μ8C, GSK2606414, PF429242 (MedChemExpress), trametinib (LC Laboratories), palbociclib (LC Laboratories), CuSO_4_ (Sigma), tunicamycin (Sigma), 4-PBA (Santa Cruz Biotechnology), cisplatin (Cayman Chemical), oxaliplatin (Cayman Chemical), doxorubicin (Sigma), salubrinal (Sigma), EGTA (Sigma), α2-3,6,8,9 neuraminidase A (NEB), LB Broth (BD Difco™), LB Agar (BD Difco™), ampicillin sodium salt (Sigma), CellEvent™ Senescence Green Flow Cytometry Assay Kit (Invitrogen), Cytofix/Cytoperm (BD Biosciences).

### Isolation of primary human and mouse NK

Human NK were isolated from discarded Leukopaks from healthy volunteer blood donors using the RosetteSep™ human NK enrichment protocol (StemCell Technologies), followed by Ficoll (GE Healthcare) density gradient centrifugation (20 min, 800g) and culture for 12-18 h in X-VIVO 10 TM medium (Lonza) supplemented with gentamicin, 5% human AB serum (Corning) and 2.5 ng/ml recombinant human IL-15 (R&D Systems). Mouse NK were isolated from splenocytes, obtained after mechanical dissociation of spleens and passage through a 40 μm sieve (BD Labware), using an NK cell magnetic purification kit (Miltenyi Biotec). Murine NK were cultured in RPMI 1640 supplemented with 10% FCS, 1% Pen/Strep and 1% L-glutamine (Gibco), and 2.5 ng/ml recombinant mouse IL-15 and 100 units of IL-2 (R&D Systems).

### Infection and tunicamycin treatment of cell lines

JEG-3 cells were infected with ZIKV for 1 h in DMEM (Gibco) with 2% FCS at an MOI of 1. Virus was aspirated and fresh culture media was added. Infected cells were cultured for indicated times before harvesting for assays. In some cases, JEG-3 were pre-treated with 10 μM salubrinal (Sigma) for 1 h before infection, and salubrinal was kept in the culture medium post infection. JEG-3 cells were infected with MOI 0.5 HSV-2 for 1 h, then medium was aspirated and cells were cultured in fresh medium for 1 or 2 d. For HCMV infection, cells were spinoculated with MOI 4 (1 h, 2800 rpm) virus, which was maintained during 12 h culture at 37° C. Spinoculation and culture were repeated once or twice to enhance infection rates and then medium was aspirated and cells were cultured in fresh medium for 12 h before harvesting. In some experiments, cells were treated with 0.5 μg/ml tunicamycin (Tu) (Sigma) for 24 h, with or without salubrinal pretreatment 1 h earlier. Salubrinal was kept in the culture throughout Tu treatment. JEG-3 viability after 1 h salubrinal pretreatment followed by 48 h ZIKV infection was measured by CellTiter-Glo Assay (Promega) following the manufacturer’s instructions. ZIKV replication under the same conditions was measured by plaque assay on Vero cells.

### Flow cytometry

For surface staining, cells were stained for 30 min on ice in the dark with LIVE/DEAD-Violet stain (1:1000) and then with primary antibodies for 15-30 min in PBS, 2% FCS (followed by secondary antibodies, when applicable, for 20 min). For protein-Ig staining, cells were incubated with 50 μg/ml of fusion protein for 1 h at 4° C and then stained with fluorescent-anti-human IgG for 1 h. Cells were fixed in 1% paraformaldehyde (Affymetrix) for 10 min before flow cytometry. For intracellular staining, cells were fixed and permeabilized using the CytoFix/CytoPerm kit. Analysis was performed on a FACSCanto II (BD). BD FACSDiva 8.0 (BD) software was used for data collection, while analysis was performed with FlowJo v10.3 (TreeStar).

### qRT-PCR

Total RNA was extracted using the RNeasy Mini kit® (Qiagen), treated with DNase (Life Technologies) and reverse transcribed using 2 μg of total RNA, 50 ng random hexamers, 400 nM dNTPs and 200 U SuperScript II reverse transcriptase (Invitrogen). Primers and diluted cDNA samples were prepared with Power SYBR Green PCR Master Mix (Applied Biosystems). Amplification cycles using the iQ5 system (BioRad) were 95° C for 10 min, then 40 cycles at 95° C for 10 sec and 55° C for 30 sec. Results were normalized to *ACTB* or *18S* RNA as indicated. Primer sequences are provided in Supplemental Table S1.

### Generation of *NCR1* (NKp46) and *NCR3* (NKp30) KO cell lines by CRISPR/Cas9

Single guide RNAs (sgRNAs) targeting the fourth exon of *NCR1* (conserved in both longest isoforms of NCR1 with transcript ID ENST00000594765) and the third exon of NCR3 (conserved in 3 of the 4 longest isoforms of *NCR3* with transcript ID ENST00000376073) were designed and cloned into the lentiCRISPR V2 vector (Addgene #52961, sgRNA sequences in Table S1). Lentiviruses were produced in HEK293T cells using standard laboratory protocols. To generate stable cell lines, YT cells (5 × 10^5^) were infected by spinoculation with viruses for each CRISPR sgRNA, using either an empty lentiCRISPR V2 or a lentiCRISPR V2 containing a non-specific sgRNA (random sgRNA) as control. Infected cells were selected (puromycin 2 μg/mL; 3 weeks) and single clones were sorted into 96-well plates using an IMI5L cell sorter (BD Biosciences). Clones were grown under selection for 3–4 wk and tested for KO by DNA sequencing and immunostaining and flow cytometry. For both *NCR1* and *NCR3*, 2 individual clones were generated from 1 or 2 independent sgRNAs for further experiments. One clone of the non-specific sgRNA (random sgRNA) was randomly chosen as control cells. To determine the allele-specific mutations, gDNA was extracted from each clone, amplified with specific primers to the target sgRNA, cloned into the CloneJET PCR cloning kit, transformed into bacteria, and at least 3 single bacterial colonies were sequenced.

### NK functional assays

NK killing of 721.221 and JEG-3, infected or not for 48 h or treated with Tu for 24 h with or without salubrinal pre-treatment or 100 μM oxaliplatin (OP) or 25 μM cisplatin (CP) for 12 h, was analyzed by 8 h ^51^Cr release assay using the indicated E:T ratios, performed in the presence of 2.5 ng/ml IL-15 (R&D Systems). NK cell killing of CRT-null HEK293T cells transfected with empty vector or WT or mutant CRT was analyzed by 4 h ^51^Cr release assay using an E:T ratio of 10:1. and YT (random sgRNA, *NCR1^−/−^*, *NCR3^−/−^*) killing of JEG-3, infected or not for 48 h, was analyzed by 8 h ^51^Cr release assay using an E:T ratio of 25:1. Murine NK killing of B16 expressing EV or WT GPI-CRT or mutated GPI-CRT or treated with OP or CP was analyzed by 4 h ^51^Cr release assay using an E:T ratio of 10:1. Murine splenic NK killing of senescent KP (treated for 6 d with trametinib (25 nM) and palbociclib (500 nM), 250 μM CuSO_4_ for 1 d or 500 μM 4-phenylbutyric acid (4-PBA) for 6 d) was measured by 8 h ^51^Cr release assay using an E:T ratio of 10:1.

To block NK receptors, NK were pretreated for 30 min at 37° C with blocking or control antibodies that were maintained during co-culture (20 μg/ml purified antibodies to NKp46 (9E2), NKp30 (P30-15), 2B4 (C1.7), NKp80 (5D12) (BioLegend), DNAM-1 (102511, R&D) or NKG2D (1D11, BioLegend) or mouse IgG1 (MOPC-21, BioLegend) as control. To block ecto-CRT, polyclonal rabbit anti-human CRT (ab2907, Abcam) was used. To measure degranulation, NK were co-cultured for 8 h with target cells in the presence of CD107a PerCP-Cy5.5 antibody (H4A3, BioLegend, 250 ng/ml) and 2.5 ng/ml IL-15 (R&D). To measure intracellular cytokine production, 2 μM monensin (BioLegend) and 3 μM brefeldin A (BD) were added after 1 h of co-culture and cells were harvested 7 h later. Cells were stained for surface markers followed by permeabilization with Cytofix/Cytoperm (BD Biosciences) and intracellular cytokine staining.

### Pulldown assay

Membrane proteins from ZIKV-infected or uninfected JEG-3 were isolated by Mem-PER Plus Kit (ThermoFisher Scientific) according to the manufacturer’s protocol. The diluted membrane fraction (300 μl in PBS) was incubated with 20 μg of NKp46-IgG1Fc or NKG2D-IgG1Fc fusion protein at 4° C with rotary agitation for 16 h and then incubated with 2 mM of DTSSP crosslinker (ThermoFisher Scientific) for 2 h on ice. The reaction was stopped with 20 mM Tris, pH 7.5. The reaction mixture was incubated with 100 μl proteinA/G-coupled Sepharose beads at 4° C with rotary agitation for 4 h. The beads were centrifuged and washed 3 times with wash buffer (10 mM Tris, pH 7.4, 1 mM EDTA, 150 mM NaCl, 1% Triton X-100, Protease inhibitor cocktail (Roche)). The beads were heated at 95° C in 100 μl 2 x SDS loading buffer and samples were analyzed by SDS-PAGE and excised bands were analyzed by mass spectrometry.

Membrane proteins from OP-treated or untreated JEG-3 or purified FC-tagged full length or truncated domains of CRT were incubated with 20 μg of NCR-Myc fusion proteins at 4° C with rotary agitation for 16 h and then with 100 μl anti-Myc coupled magnetic beads (Genscript) at 4° C with rotary agitation for 4 h. The beads were magnetically separated and washed 3X with wash buffer. The beads were then heated at 95° C in 100 μl 2 x SDS loading buffer and the samples analyzed by SDS-PAGE and immunoblot.

### Phospho-flow cytometry analysis of NKp46 signaling

JEG-3 (5×10^6^), pre-treated with OP for 4 h at 37°C and with 10 μg/ml of anti-CRT or isotype control antibody for 30 min, were incubated for 15 min at 37°C with WT or *NCR1^−/−^* YT (10^6^), pretreated for 30 min with 10 μg/ml of anti-NKG2D. Cells were then fixed with Phosflow Fix Buffer I and permeabilized with Phosflow Perm Buffer III (BD Bioscience) and stained for CD56, p-CD3ζ, p-Syk, and p-Tyr before flow cytometry analysis on gated CD56^+^ cells.

### Protein expression and purification

The full-length or truncated N, P, C domain coding sequences of human CRT were cloned into the pCMV vector (Addgene plasmid #59314) to generate recombinant constructs with C-terminal Human IgG1Fc (Ig) tags. The extracellular domain of NKp46 was cloned into pcDNA Myc-His 3.1a vector (Invitrogen) or pCMV vector (Addgene plasmid #59314) to generate a recombinant construct with a C-terminal Myc or Human IgG1Fc (Ig) tag. All plasmids were verified by DNA sequencing and transfected into HEK293T. Recombinant proteins were purified as previously described^55^. cDNA clones of human *CALR* and *NCR1* were obtained from the DF/HCC DNA Resource Core at HMS and primers used are given in Table S1.

### Microscale Thermophoresis (MST)

Purified human MYC- or Fc-tagged NKp46 were labeled with Alexa647 using the microscale protein labeling kit (ThermoFisher Scientific). Alexa647-labeled NKp46-Myc (50 nM) in MST buffer (50 mM HEPES, 150 mM NaCl, 0.05% Tween20) was mixed with 0-100 μM Fc-tagged full-length or N, P, or C domains of CRT or other recombinant His-tagged ligands and incubated at room temperature for 30 min to achieve binding equilibrium. The reaction mixtures were taken up into MST capillaries and measurements were acquired using a Monolith NT.115 (NanoTemper Technologies) microscale thermophoresis instrument. The data was fit using the Hill equation and K_d_ values were determined using MO.Affinity Analysis software (NanoTemper Technologies, Munich, Germany). Sialic acid was removed from NKp46 protein by treatment with neuraminidase A for 1 h at 37°C.

### Raman Spectroscopy

Raman spectroscopy was carried out using an Horiba XploRa confocal Raman microscope. Spectra were collected using a 785 nm laser (~41 mW) at 1 cm^−1^ resolution. The spectrometer slit was set to 200 μm with 500 μm aperture. 20 μM NKp46 was mixed with 5-20 μM CRT in sodium phosphate buffer in 20 μl. Acquisition was carried out at room temperature in triplicate samples. Spectral readouts were normalized to the sharp peak at 330 cm^−1^. The fluorescent baseline was fit using a polynomial fit and subtracted before peak deconvolution was perfomed in Labspec 6. The change in the peak area at ~1658 cm^−1^ (obtained after Lorentzian fitting in the AmideI deconvolution step for each acquisition) was plotted against CRT concentration. The peak at 1658 cm^−1^ was chosen as the marker for the interaction in the Raman spectrum of the amide I region since it was found to be a characteristic of the NKp46-CRT interaction. A hyperbolic model was used to calculate Kd value. The quality of the model used to fit the data was assessed by calculating the R2 value, which was 0.907.

### Molecular modeling and docking

Using NKp46 and CRT sequences, model structures were generated using the threading technique by the stand-alone version of I-TASSER. Energy minimization and other subsequent processing were carried out using GROMACS and Chiron before initiating molecular docking. Molecular docking was performed using ClusPro program, which works on an ensemble pipeline involving FFT (fast Fourier transform)-based global sampling and the PIPER algorithm followed by a clustering technique to find highly populated clusters of lowest energy and then CHARM energy minimization to avoid steric clashes. Docked complexes were further subjected to energy minimization to remove steric clashes with Chiron. The docked complex selected for subsequent analysis was screened using lowest cluster-energy principles. The interaction energy between two proteins in PIPER was calculated using the formula E=0.40E_rep_+(−0.40E_att_)+600E_elec_+1.00EDARS. E_rep_ and E_att_ represent the repulsive and attractive contributions to the van der Waals interaction energy. E_elec_ denotes an electrostatic energy term while EDARS denotes pairwise structure-based potential and primarily represents desolvation contributions, i.e., the free energy change due to the removal of water molecules from the interface. To denote the possible contributions of charged residues in stabilising the NKp46-CRT interaction, surface charges in the bound complexes were rendered using the Vacuum electrostatics mode of PyMol, where surface charges are represented by a red (negative) to blue (positive) scale.

### Generation of *CALR^−/−^* HEK293T

CRT-null HEK293T cells were generated using Guide-it CRISPR/Cas9 System (Takara) according to the manufacturer’s protocol. Briefly, two guides were designed targeting exon2 (sgRNA1:GCCGTCTACTTCAAGGAGCA, sgRNA2: TAATCCCCCACTTAGACGGG). The guides were chosen to minimize off-target effects using (http://crispr.mit.edu/). Plasmids expressing individual guides were transfected into HEK293T by Amaxa nucleofector and Nucleofector® Kit V (Lonza). After 48 h, green fluorescent cells were sorted using a BD Aria cell sorter. Two sequential rounds of transfection were carried out and single clones were sorted in 96 well plates. Two single clones, selected based on immunoblot analysis, were verified by sequencing.

### Site Directed Mutagenesis and Complementation

Point or deletion mutations were prepared from human or mouse CRT expression clones in pCMV3-C-Myc (SinoBiological) using the Q5® Site-Directed Mutagenesis Kit (NEB) according to the manufacturer’s protocol and primers designed using the NEBaseChanger™ web tool. Plasmid sequences were validated by sequencing. Complementation of CRT-null cells was performed by lipid transfection of mutated or WT plasmids using Lipofectamine 2000 (Invitrogen) in 6 well plates. Wells with >85% transfected cells were analyzed by flow cytometry for cell surface CRT and NKp46-Ig binding and used for NK killing assays.

### Imaging flow cytometry

B16 (EV or GPI-CRT or treated with either CP or OP) were stained with CellTracker Red CMTPX (CMTPX) (ThermoFisher Scientific) and cocultured with WT or *Ncr1*^−/−^ mouse NK at an E:T ratio of 1:5 for 30 min. Cells were fixed with 4% paraformaldehyde (Sigma) for 10 min and stained with rabbit polyclonal anti-CRT (MBL Bioscience), rat monoclonal anti-NKp46 PE-Cy7 (BioLegend), and rat monoclonal anti-CD49b-APC (BioLegend) before analysis on an ImageStream Amnis X MKII (Amnis) using Ideas software (Amnis). Cell doublets (conjugates) were selected based on aspect ratio versus cell area, gating on CMTPX^+^CD49b^+^ (tumor/NK) conjugates.

### Mouse studies

C57BL/6 and B6;129-Ncr1tm1Oman/J (*Ncr1^−/−^)* female mice (6–8 weeks old) were purchased from The Jackson Laboratory. Animal use was approved by the Animal Care and Use Committees of Boston Children’s Hospital and Harvard Medical School. For tumor challenge experiments, B16 EV or GPI-CRT-expressing cells (2×10^5^) were injected subcutaneously into the right flank of WT (n = 7) and *Ncr1^−/−^* C57BL/6 (KO) (n = 7) mice and tumor volumes were measured every other day. For the DOX experiment, B16 (2×10^5^) were injected subcutaneously into the right flank of WT (n = 7) and *Ncr1^−/−^* C57BL/6 (n = 7) mice and animals were injected iv 6 and 13 day later with 5 mg/kg DOX. Tumor growth was monitored by measuring the perpendicular diameters of tumors every other day. When control tumors grew to ~1000 mm^3^ volume, all mice were sacrificed and tumors were collected for analysis. In the experimental metastasis assay, B16 EV or GPI-CRT-expressing cells (2.5×10^5^) were injected into the tail vein of WT or *Ncr1^−/−^* mice and were monitored thereafter every other day. After 21 or 28 days, mice were euthanized and numbers of tumor colonies on lung surfaces were counted using a dissecting microscope (OLYMPUS).

In some experiments, WT mice were injected ip daily for 2 d before tumor implantation with antibodies to CD8^+^ T cells (0.5 mg/mouse rat anti–mouse CD8 mAb, clone 2.43, BioXCell), NK cells (0.2 mg/mouse of rabbit anti–mouse NK1.1, BioXCell) or 300 μg/per mouse anti-CSF1R (BioXCell) or a control antibody (LTF-2, BioXCell). Thereafter, for the duration of the experiment, CD8 and NK1.1 antibodies were injected once a week and CSF1R every other day. Cell depletion was verified by flow cytometry analysis of blood cells at the time of tumor implantation and at the time of sacrifice.

For the KP cancer model, 2×10^5^ KP cells in 100 μl PBS were injected sc in the right flank of WT, *Ncr1^−/−^* or *Klrk1^−/−^* C57BL/6 mice (n = 5/gp). Mice were monitored for tumor growth by caliper measurements every other day. Mice, randomly assigned to study groups 14 d after implantation, were treated with vehicle or trametinib (1 mg/kg) and palbociclib (100 mg/kg) for 5 consecutive days. When control tumors grew to ~1000 mm^3^, all mice were sacrificed and tumors were collected for analysis.

Whole tumors were collected, cut into small pieces and treated with 2 mg ml^−1^ collagenase D, 100 μg ml^−1^ DNase I (both from Sigma) and 2% FBS in RPMI with agitation for 30 min. Tumor fragments were homogenized and filtered through 70-μm strainers to obtain single cell suspensions and then stained and analyzed by flow cytometry gating on CD45^+^ cells. Isolated cells were stained with CD45-PerCPCy5.5 (clone 30-F11), CD8-PacBlue, -PerCPCy5.5, -Alexa700, -fluorescein isothiocyanate (FITC) or -APC (clone 53-6.7), CD4-PE-Cy7, -APC or -PerCPCy5.5 (clone GK1.5), CD49b-FITC, -PacBlue or -PerCPCy5.5 (clone DX5), NKp46-APC (clone 29A1.4), CD11b-Alexa700 (clone M1170), F4/80-PE-Cy7 (clone BM8) and CD44-PerCPCy5.5 (clone IM7) (BioLegend). Dead cells were excluded using the Zombie Yellow Fixable Viability dye (BioLegend). CD45^+^ lymphocyte populations were further defined as: CD8^+^ T cells (CD3^+^CD8^+^), CD4^+^ T cells (CD3^+^CD4^+^), NK cells (CD3^−^CD49b^+^), dendritic cells (F4/80^−^CD11c^+^), tumor-associated macrophages TAM (F4/80^+^CD11b^+^). For intracellular staining, cells were first stained with antibodies to cell-surface markers for 30 min, then fixed and permeabilized with fixation/permeabilization buffer (BD Pharmingen) and stained with GzmB-PacBlue (clone GB11, ThermoFisher Scientific), perforin-PE (clone S16009B, BioLegend). For intracellular cytokine staining (ICS), 10^6^ cells per sample were cultured in RPMI containing 2% FBS and stimulated with PMA (50 ng ml^−1^, Sigma), ionomycin (2 μg ml^−1^, Sigma) and Golgi-plug (1.5 μl ml^−1^, ThermoFisher Scientific) for 4 h. Cells were then stained with IFNγ-PacBlue (clone XMG1.2) and TNF-phycoerythrin-Cy7 (clone MP6-XT22) after fixation/permeabilization.

### Senescence Induction

A549 or KP were pretreated or not for 6 d with trametinib (25 nM) and palbociclib (500 nM), 250 μM CuSO_4_ for 1 d or 500 μM 4-phenylbutyric acid (4-PBA) for 6 d and washed 3 times with PBS before use. Cells were stained for ICAM-1, MICA/B, CRT or RAE1 for flow cytometric analysis. SA-β-galactosidase activity was measured using the CellEvent Senescence Green Flow Cytometry Assay Kit (ThermoFisher Scientific) following the manufacturer’s protocol. To assess killing, cells were co-cultured with NK in the presence or absence of indicated blocking antibody at an E:T ratio of 25:1 for 8 h.

### Statistical analysis

Statistical analysis was performed using GraphPad Prism V9.1.2. Prior to applying statistical methods, whether the data fit a normal distribution was evaluated by the D’Agostino and Pearson normality test. The distribution was considered normal when *P* ≤ 0.05. Parametric or non-parametric (Mann-Whitney test) two-tailed unpaired *t*-tests were used to compare two groups. Column multiparametric comparison were analyzed by one-way ANOVA using the Kruskal-Wallis test and Sidak’s or Tukey’s multiple comparisons test. Two-tailed *Chi*-square test was used for categorical column statistics in Fig. 3g,h. Multiple groups were compared by two-way ANOVA with additional Tukey’s multiple comparisons test. Tumor growth curves were analyzed by first calculating the area under the curve, followed by two-tailed non-parametric unpaired t-test. Killing assays employing different E:T ratios were analyzed by area under the curve, followed by non-parametric one-way ANOVA with Tukey’s multiple comparisons test. Non-parametric tests were used in experiments with animal samples because of the non-normal distribution of these data. Differences were considered statistically significant when *P* ≤ 0.05. Most experiments were not blinded as to group allocation, either while collecting data or assessing the results.

## Data Availability

The data that support the findings of this study are available from the corresponding authors upon request.

Further information on research design is available in the Nature Research Reporting Summary linked to this article.

## Data reporting

No statistical methods were used to predetermine sample size. The experiments were not randomized and the investigators were not blinded to outcome assessment.

## Acknowledgments

This work was supported by US NIH grant AI150671 (J.L.) and a Jeffrey Modell Foundation fellowship (S.S.S.) and internship (M.R.). We thank Ofer Mandelboim (Hebrew University Faculty of Medicine) for NKp46-Ig protein, Tyler Jacks (Koch Institute, MIT) for the KP cell line, and Justin Kline (Department of Medicine, University of Chicago) for pRetroX-IRES-GPI-CRT-DsRed plasmid.

## Author Contributions

S.S.S. and J.L. conceived the study; S.S.S., C.J. and J.L. designed the experiments and analyzed the data; S.S.S., A.C., C.J, D.L., J.H, K.M., Y.Z., M.R., and S.C. performed the experiments; H.W. and J.L. supervised the experiments and data analysis; and S.S.S. and J.L. wrote the manuscript.

## Author Information

The authors declare no competing financial interests. Correspondence and requests for materials should be addressed to J.L.

